# Segment number threshold determines juvenile onset of germline cluster proliferation in *Platynereis dumerilii*

**DOI:** 10.1101/2021.04.22.439825

**Authors:** Emily Kuehn, David S. Clausen, Ryan W. Null, Bria M. Metzger, Amy D. Willis, B. Duygu Özpolat

## Abstract

Many animals rely on sexual reproduction to propagate by using gametes (oocytes and sperm). Development of sexual characters and generation of gametes are tightly coupled with the growth of an organism. *Platynereis dumerilii* is a marine segmented worm which has been used to study germline development and gametogenesis. *Platynereis* has 4 Primordial Germ Cells (PGCs) that arise early in development and these cells are thought to give rise to germ cell clusters found across the body in the juvenile worms. The germ cell clusters eventually form the gametes. The stages of germline development and how the 4 PGCs become the numerous germ cell clusters are not well-documented in the juvenile stages. *Platynereis*, like other segmented worms, grows by adding new segments at its posterior end. The number of segments generally reflect the growth state of the worms and therefore is a useful and easily measurable growth state metric. To understand how growth correlates with development and gametogenesis, we investigated germline development across several developmental stages using germline/multipotency markers. We found that segment number predicted the state of germline development and the abundance of germline clusters. Additionally, we found that keeping worms short in segment number via changing external conditions or via amputations supported segment number threshold requirement for germline development. Finally, we asked if these clusters in *Platynereis* play a role in regeneration (as similar free-roaming cells are observed in *Hydra* and planarian regeneration) and found that the clusters were not required for regeneration in *Platynereis*, suggesting a strictly germline nature. Overall, these molecular analyses suggest a previously unidentified developmental transition dependent on the growth state in juvenile *Platynereis* when germline proliferation is substantially increased.

## INTRODUCTION

Sexual maturation is one of the major developmental transitions in an animal’s life history. Maturation requires production of gametes (eggs and sperm) via a process called gametogenesis. The onset and progression of gametogenesis are tightly coupled with the growth and nutritional state of an individual. Therefore growth, nutritional state, and sexual reproduction are intertwined [1]. Animals have mechanisms that assess growth and regulate processes such as gametogenesis as a result of the systemic check and feedback [2,3]. For example, in *Drosophila* and *C. elegans*, nutritional status and insulin signals affect germline stem cell (GSC) division rate and gamete production [4–6]. In insects, reaching a critical weight is required for the transformation from an immature form to an adult (metamorphosis) which will reproduce sexually [7]. This somatic-germline crosstalk and the regulation of maturation based on body weight and size has been extensively studied in many insects and other ecdysozoans, which have stepwise (discontinuous) growth with molting and/or metamorphosis [8,9]. However, there is much less information available on how growth and nutritional status affects developmental transitions in invertebrates which display continuous growth, such as annelids (segmented worms), planarians, and tunicates.

Annelids have repeating body parts called segments. Most annelids grow continuously by adding segments at their tail end throughout their lives. As a result, growth can be easily measured by counting the number of chaetigerous segments (segments that have bristles, aka. setigers) and correlations can be made with developmental transitions and stages of sexual maturation. We use the polychaete *Platynereis dumerilii* as a model system for studying germline formation, gametogenesis, and germline regeneration. *Platynereis* has 4 primordial germ cells (PGCs) that are set aside during embryogenesis [10–14]. The PGCs are quiescent until they migrate anteriorly (Fig. 1A-B) where they start proliferating and form a cluster (anterior cluster, Fig. 1C). They express germline markers such as *vasa*, *piwi*, *nanos*, and express the Vasa protein (Fig. 1D’). As the worms grow, this anterior cluster located right behind the jaws (around segments 4-5 in younger juveniles) is thought to give rise to cells that migrate posteriorly to the trunk segments and form many small gonial clusters spread across the coelomic cavity, in the parapodia (appendages on each side of a segment) and around the gut [13] (Fig. 1D-D’). These clusters eventually give rise to the gametes [15–17] (Fig. 1E). Overall, the behavior of 4 PGCs in the early larval stages, and late stages of gametogenesis in the older juveniles have been well-documented (Fig. 1A-B and Fig. 1E), there is less documentation for the earlier gametogenesis stages, namely the process of numerous posterior small *vasa*+ gonial cluster formation during the mid-juvenile stages (Fig. 1C-D).

**Figure 1.**
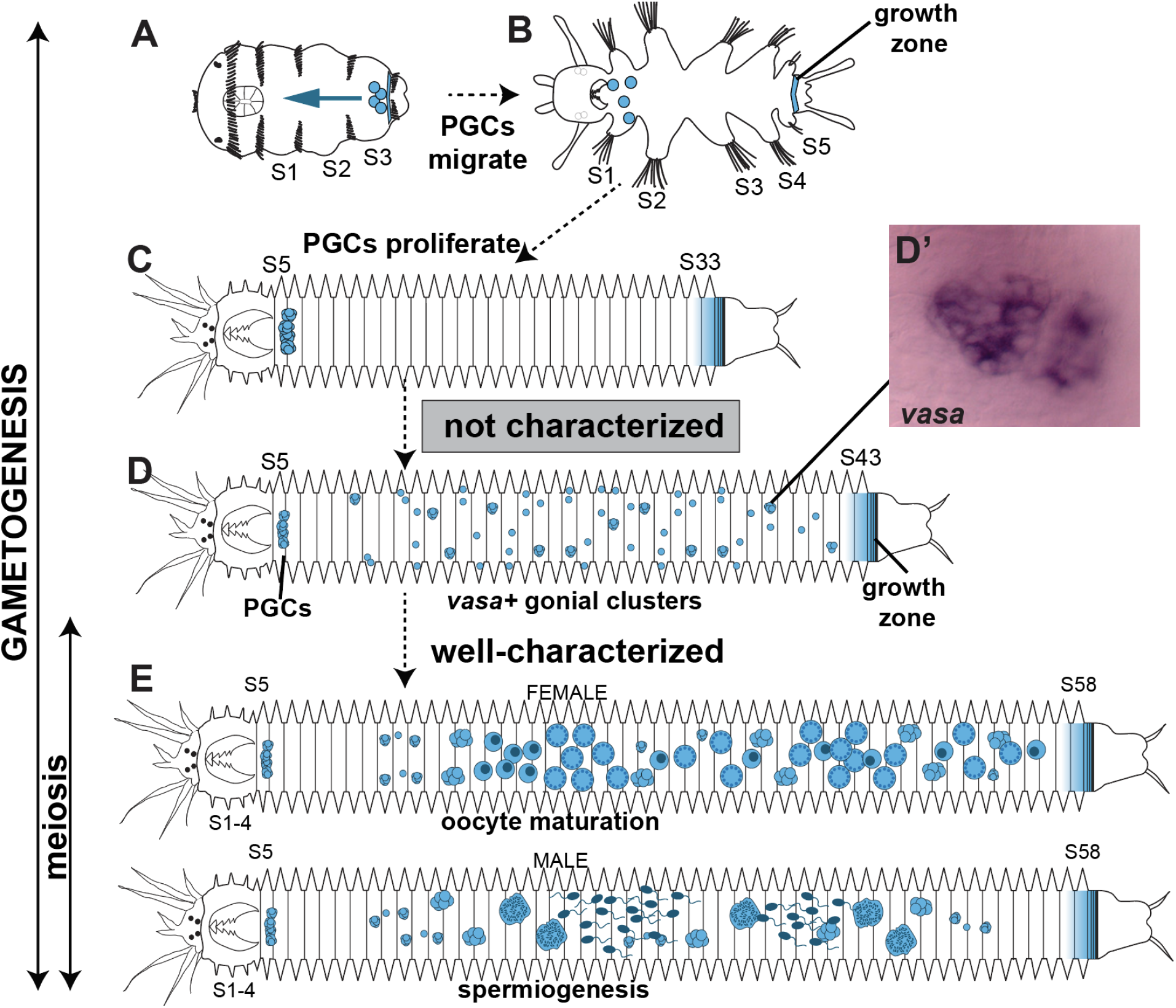
Knowledge gap in *Platynereis* gametogenesis. **A)** 3-segmented larval *Platynereis* has 4 PGCs (4 blue circles) located next to the growth zone at the tail. **B)** By the time the larva grows into 5 segments, 4 PGCs migrate anteriorly. **C)** In juvenile worms these cells proliferate and form a large cluster in segments 5-6. **D)** Many small clusters that later appear in posterior segments are thought to come from the large anterior cluster but there is no direct evidence. **E)** Later stages of gametogenesis where sperm and oocytes are easier to distinguish have been well-documented. S: segment

Gametogenesis in many animals comprises a wide range of events: first, PGCs proliferate via mitosis to increase their numbers and produce gonial cells (spermatogonia and oogonia). These gonial cells may continue to multiply via mitosis and eventually form primary oocytes and primary spermatocytes. At this stage meiosis starts which becomes evident by nuclear changes. In parallel, other gamete maturation processes such as vitellogenesis in oocytes take place, and eventually the mature oocytes and sperm are produced [18,19]. In *P. dumerilii*, detailed work has been carried out on the late stages of gametogenesis using TEM and other histological techniques [15–17,20,21]. These studies investigated worms that are 50 segments or longer, in which gamete maturation has already begun. However, coelomic gonial clusters at the earlier stages of gametogenesis are hard to reliably observe and differentiate from somatic cells without molecular markers. Because the early works predated the availability of molecular tools and markers, the early stages of gametogenesis when gonial clusters form and populate the trunk segments in the immature juveniles were not investigated. More recently molecular gene expression analyses have been carried out in *P. dumerilii* using several germline/multipotency markers [11–13]. These workers hinted at a connection between segment number and changes in gonial cluster distribution but they have not systematically investigated juveniles of different segment lengths. Overall, at what stage gonial clusters start populating the trunk segments, and whether there is a critical size for this step of gametogenesis to start have not been identified.

In this study, in order to address this knowledge gap, we analyzed the dynamic behavior of *vasa*-expressing gonial clusters across developmental stages of the juvenile *P. dumerilii*. We found that the formation of *vasa*+ gonial clusters consistently occurs after worms have reached 35-40 chaetigerous segments in length. We show that the chronological age of the worms is a poor predictor of the number of gonial clusters a worm would have. This indicates that gametogenesis in *P. dumerilii* is tightly coupled with growth and not age, and there appears to be a critical size (in terms of segment number) that needs to be reached for progression of gametogenesis. This molecular and quantitative approach allowed us to determine a developmental stage transition that was not identified before. We propose the 35-40 segment growth phase to be the onset of sexual maturation in this species. These analyses will facilitate further investigations of gametogenesis as well as germline regeneration, and will help clarify the relationship between growth and sexual maturity in a semelparous organism with indeterminate growth.

## MATERIALS AND METHODS

### GitHub resources for this publication

We have uploaded R code, associated spreadsheets, protocols, and additional resources related to the work reported here on GitHub to a specific repository for this manuscript [22].

### Animal culturing

Animals were raised in conditions as described previously [23] except where noted below. Briefly, all the samples used in this study were kept at Light/Dark photoperiodic conditions of 16:8 hours, and 8 days of moonlight at night every 28 days.

### Whole mount *in situ* hybridizations (WISH)

We followed the published WISH protocols with some modifications [22,24]. Briefly, animals were fixed in 4% paraformaldehyde at 4°C overnight, transferred stepwise into 100% methanol, and kept at −20℃ until the start of WISH procedure. Samples were rehydrated through a methanol/DEPC-treated PBSt series. They were then digested in 0.1 mg/mL Proteinase K (Ambion AM2548) for 5 minutes at room temperature before being briefly (<1 minute) washed in a glycine (2mg/mL) DEPC-PBSt solution, and refixed in 4% paraformaldehyde for 30 minutes on ice. Dig-labeled sense and antisense RNA probes for *vasa* were prepared as described previously [24] (Ambion AM1310, Roche 11277073910). Probes were diluted to 2ng/μL in 100% hybridization solution, and were denatured at 85℃ for 10 minutes. Samples were then incubated in the probe solution at 65℃ overnight (approximately 16 hours). Final colorimetric reaction was developed using NBT/BCIP substrate. Keeping most samples in NBT/BCIP overnight typically provided enough time for the signal to develop with minimal background. Samples were washed twice with stop buffer to end the reaction and at least three times in PBSt (30 minutes each) while being kept on ice and on a nutator. Samples were then transferred into 75% glycerol/25% PBSt stepwise and stored (protected from light) at −20℃ until imaging. Sense probe controls were included in each experiment to account for background staining.

### Hybridization Chain Reaction (HCR)

#### Probe (oligo pool) design

Based on the split-probe design of HCR3.0 reported previously [25,26], we wrote a custom software [27] to create 30 DNA oligo probe pairs specific to *Platynereis dumerilii vasa* mRNA (Genbank Accession: AM114778.1). For each probe pair, the software takes a 52bp footprint from the input cDNA sequence, using the 25 nucleotides at the 5’ and 3’ ends as the anti-sense complement of the RNA target sequence, the intervening 2bp is left unpaired as a buffer region between the pair. The algorithm then skips ≥2bp 3’ of the previous probe pair and repeats the creation of a new probe pair. Finally, one half of the B1 initiator (B1I1 or B1I2) is appended to each oligo in a pair, such that each pair has a complete B1 initiator sequence between the two of them. Exact oligo sequences and the associated hairpins are listed in the Supplementary Materials file. The sequences generated by the software were used to order a single, batched DNA oligo pool (50μmol DNA oPools Oligo Pool) from Integrated DNA Technologies, resuspended to 1μmol/μl in 50mM Tris buffer, pH 7.5.

#### Tissue collection and HCR

Samples for HCR were fixed the same way as the WISH samples explained above and stored in 100% Methanol at −20C until the procedure. Probe hybridization buffer, probe wash buffer, amplification buffers, and a DNA HCR amplifier hairpin set were purchased from Molecular Instruments (https://www.molecularinstruments.com/). The sequence for the HCR B1 amplifier has been reported previously [25] (see Suppl. Material). HCR was performed according to the published methodology [26]. To suit our samples, steps from tissue rehydration through to post-fixation were performed in 6 well plates according to standard *Platynereis* colorimetric *in situ* protocols [22]. All subsequent steps were performed in 2mL tubes. The wash volume was decreased from 2mL to 800uL to accommodate less tissue and conserve reagents. Incubation volumes were maintained. At the amplification step, DAPI was added to a final concentration of 2ug/mL.

#### Imaging

Samples were mounted in Slowfade Glass with DAPI (Thermo S36920-5X2ML), kept at 4C until imaging, and imaged using Zeiss Laser Scanning Confocal Microscope LSM 780. Images were processed for levels in FIJI.

### Scoring control samples for *vasa*+ clusters

We culture worms in plastic boxes labeled with the date of birth of a given batch, therefore the chronological age of any worm in our cultures is known at the time of fixation. We scored each worm sample for the total number of chaetigerous segments and the total number of *vasa*+ clusters. For assembling this dataset, we collected and fixed worms at a range of chronological ages and a range of segment numbers, then processed and analyzed them for the *vasa* expression phenotypes. We have also included control worms from other experiments into this dataset. More than 400 control worms were scored.

For scoring first, samples were imaged using a Zeiss V20 stereoscope and scoring was carried out on the images. Then, the number of segments and *vasa*+ clusters on each sample was counted using the count tool in Adobe Photoshop or the cell counter plug-in in FIJI [28]. Examples have been uploaded to Zenodo [22] (doi.org/10.5281/zenodo.4587817, doi.org/10.5281/zenodo.4587804). The following criteria were used for scoring: i) *vasa*+ gonial clusters were identified by their distinctive dark staining and well-defined borders. However, to avoid counting false positives, clusters that looked to be located in parts of the worm containing large amounts of gut content were not counted. Likewise, clusters located in the parapodia that displayed similarity to the background signal in the sense controls were not counted. ii) the anterior cluster and any clusters near the head segments that looked like could be a part of the anterior cluster (~first 6 segments) were not included. In all the scatter plots, the cluster numbers exclude the anterior cluster. iii) Positive clusters were identified as strongly stained single cells or globular clusters of multiple cells. For simplicity, we refer to anything that was counted as positive for the *vasa*+ marker and received a scoring pointer as a “cluster”. Although some clusters consisted of multiple cells, it was not possible to count the exact number of cells in each cluster. If clusters appeared to be touching but the overall shape indicates that there were at least two clusters, these were counted as two separate clusters. iv) For counting segments, presence of chaetae was deemed sufficient to count a segment, including for the young segments at the posterior end. Each bilateral pair of parapodia was considered to be one segment. v) During dehydration-rehydration and WISH, sometimes the tails broke. Any worms with tails that had broken off during the WISH procedure or tails that looked like they were regenerating in the control group were also removed from the data pool to prevent uncertainty about the number of segments a broken sample had.

### High vs. low populations density experiments

A fertilization batch consisting of full and half-sibling worms was kept in a large culture box (35.6 cm × 27.9 cm × 8.3 cm, *Sterilite*, 1963) from the time of fertilization until 6 weeks post fertilization. At this time, four low density and four high density cultures were established by transferring worms from the original culture into separate small boxes (27.9 cm × 16.8 cm × 7 cm, *Sterilite*, 1961) in 500 mL NFSW. High density cultures consisted of 100 worms per box while low density cultures consisted of 20 worms per box. Half of all low- and high-density boxes received the normal amount of food per worm (0.57 mg of spirulina/sera micron mixture per worm) each time they were fed. Remaining cultures received half of the normal amount of food (0.28 mg). All culture boxes were fed three times per week (Monday, Thursday, and Saturday). Before each feeding, the seawater in each box was changed to prevent fouling. A whole box from each group (high or low) and feeding regimen (normal or underfed) was collected and fixed for WISH after two and four weeks in these cultures. Worms from the original culture were fixed at the time the new high- and low-density boxes were established (t=0).

### P10-P20 Amputation Experiment

Prior to amputation, worms were anesthetized in a 1:1 mixture of 0.22 μm natural filtered seawater (NFSW) and 7% MgCl2 until no movement was observed when prodded (approximately 5-10 minutes). The segment number was determined by counting lateral pairs of parapodia. Amputations were made after the 10th or 20th segment by making a transverse cut using a scalpel (Dynarex 4115). Worms were washed in 0.22 μm NSFW and placed into clean small (27.9 cm × 16.8 cm × 7 cm, *Sterilite*, 1961) culture boxes. The samples were fixed at the following time points: day 0 (no amputation), 3 days post-amputation (dpa), 5 dpa, 7 dpa, 14 dpa, 20 dpa, 25 dpa, 30 dpa, 45 dpa, and 60 dpa. Among these time points, 14 dpa and older were used for the P10-P20 amputation analyses. Examples from all the time points, (including earlier ones than 14 dpa) were used for the regeneration analyses.

### Plots and Statistical Analyses

#### Scatter plots and boxplots

Scatter plots and boxplots were generated either in RStudio [29,30] or Microsoft Excel. The visuals were amalgamated and edited in Adobe Illustrator for clarity, shape, and text size adjustments. All R scripts and datasets (01, 02, 03) are available as Supplementary Material via GitHub [22].

#### Regression analyses of control samples for *vasa*+ clusters (Dataset-01)

To estimate the conditional median number of *vasa*+ clusters at each observed number of segments, we fit an isotonic median regression via the gpava function in the isotone R package [31]. In our analysis, we excluded observations for which worms broke prior to measurement of vasa+ cluster and segment number, as well as two observations for which age and/or posterior vasa+ cluster count was missing. We constructed 95% pointwise confidence intervals for the conditional median via a cluster bootstrap. The clusters were defined by the culture boxes in which each worm was grown in order to account for within-culture box dependence. We used 10,000 bootstrap iterations, and applied the correction given in Table 1 by Abrevaya et al [32].

#### Regression analyses of P10-P20 amputation experiment (Dataset-03)

In this experiment, the relationship between segment number and conditional median number of *vasa*+ clusters were compared across three conditions: control condition, worms after the 10th segment (P10) and worms amputated after the 20th segment (P20). We excluded all observations on worms with broken tails (n = 150). While the probability of tail break is likely associated with the number of segments, we analyzed complete worms under the assumption that for worms of a given length, the number of *vasa*+ clusters is not associated with the probability of tail break. If this assumption holds, this missing data reduces our sample size (n = 353, after exclusions) but should not bias our estimates of the association between number of segments and conditional median number of *vasa*+ clusters. For each of these conditions, we estimated conditional median *vasa*+ clusters via the gpava function in R package isotone and constructed 95% pointwise confidence intervals as described above, again with cluster resampling based on culture box.

## RESULTS

### Gametogenesis in the juvenile and mature *Platynereis*

To characterize the behavior of the *vasa*+ PGCs and gonial clusters across growth stages of juvenile *P. dumerilii*, we carried out whole mount *in situ* hybridization for *vasa* (Fig. 2). We found that juvenile worms that are around 10-19 segments long (n=30) only had a few cells (4 or more) arranged in an arch right behind the head region (Fig. 2A-A’). These cells appeared to multiply as the worms reached a length of 20-29 segments (n=110), forming a ring-like structure in many of the samples (Fig. 2B-B’). Neither of these stages (Figs. 2A-B) had any *vasa*-expressing cells in the other segments except in the arch-like formation behind the head at segments 4-6 and at the posterior growth zone (Fig. 2C’’). In the worms that were 30-39 segments long (n=76), the ring structure turned into a broader band (Fig. 2C-C’). In this group, worms longer than 35 segments started having few cells expressing *vasa* in the trunk region (Fig. 4B). This pattern changed drastically as the worms reached lengths longer than 40 segments (n=110): in most of the worms longer than 40 segments, we detected numerous *vasa*+ cells and cell clusters scattered across the body (Fig. 2D-F’, Fig. 4B, Suppl. Fig. 1). In worms longer than 50 segments (n=168), we observed a few different states. Many worms in this group had regularly spaced gonial clusters, especially in more anterior trunk segments (Fig. 2F-F’). Meanwhile, in some samples we observed oocytes at different stages of maturation and expressing *vasa* (Fig. 2H-H’), as well as samples that had *vasa* expression only in the parapodia (Fig. 2G-G’). We think these may be males starting to make sperm and getting closer to completion of maturation. Sense controls showed an occasional striated background staining (Suppl. Fig. 4A’). Also note that the chaetal sacs in parapodia have a characteristic background staining in most juveniles usually in the more posterior segments (Suppl. Fig. 4A’’). Overall, *vasa*+ gonial clusters in the trunk segments started appearing in worms 35 segments or longer, they became more prevalent once worms surpassed 40 segments in length, and in some worms longer than 60 segments maturing gametes were evident.

**Figure 2.**
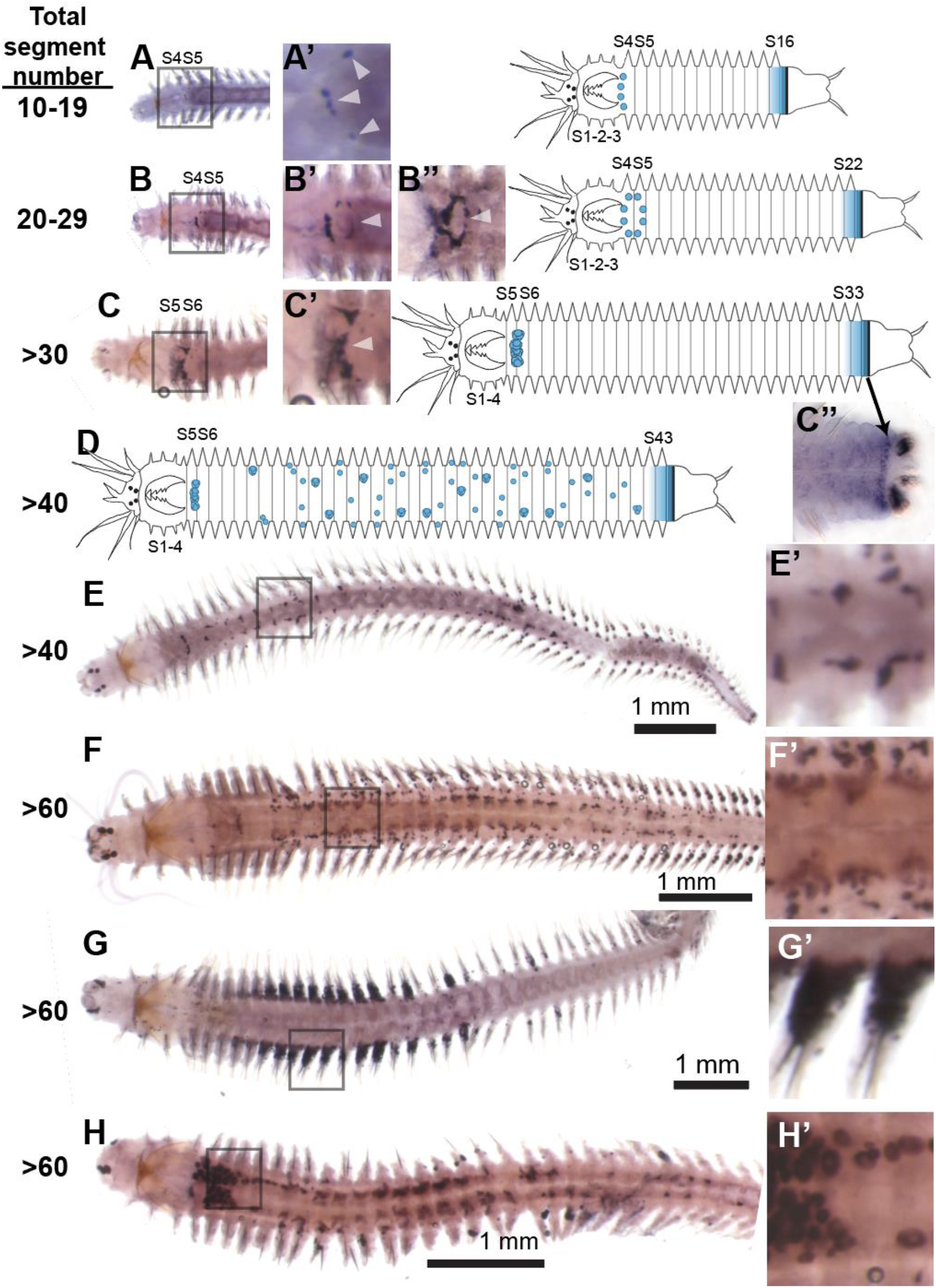
Gametogenesis in the immature juvenile and premature *Platynereis*. A) In worms between 10-19 segments in length, vasa positive cells can be found towards the anterior of the worm, just behind the jaws. B) More vasa positive cells appear in a ring-like structure to encircle the esophagus as the worm grows. C) Vasa expression continues to accumulate in this region in worms 30-39 segments long. D-E) Clusters of vasa positive cells begin to appear throughout the posterior body cavity of the worm once it has reached 40 segments in length. F) Clusters are more numerous in larger worms. G-G’) Vasa signal becomes localized in the anterior parapodia of some worms. H) While many worms do not display any vasa clusters between segments 7-10, we have seen a few exceptions. It is unclear whether vasa positive clusters proliferate in the anterior of the worm and then migrate posteriorly, or if vasa expression originates regardless of cell lineage.

**Figure 3.**
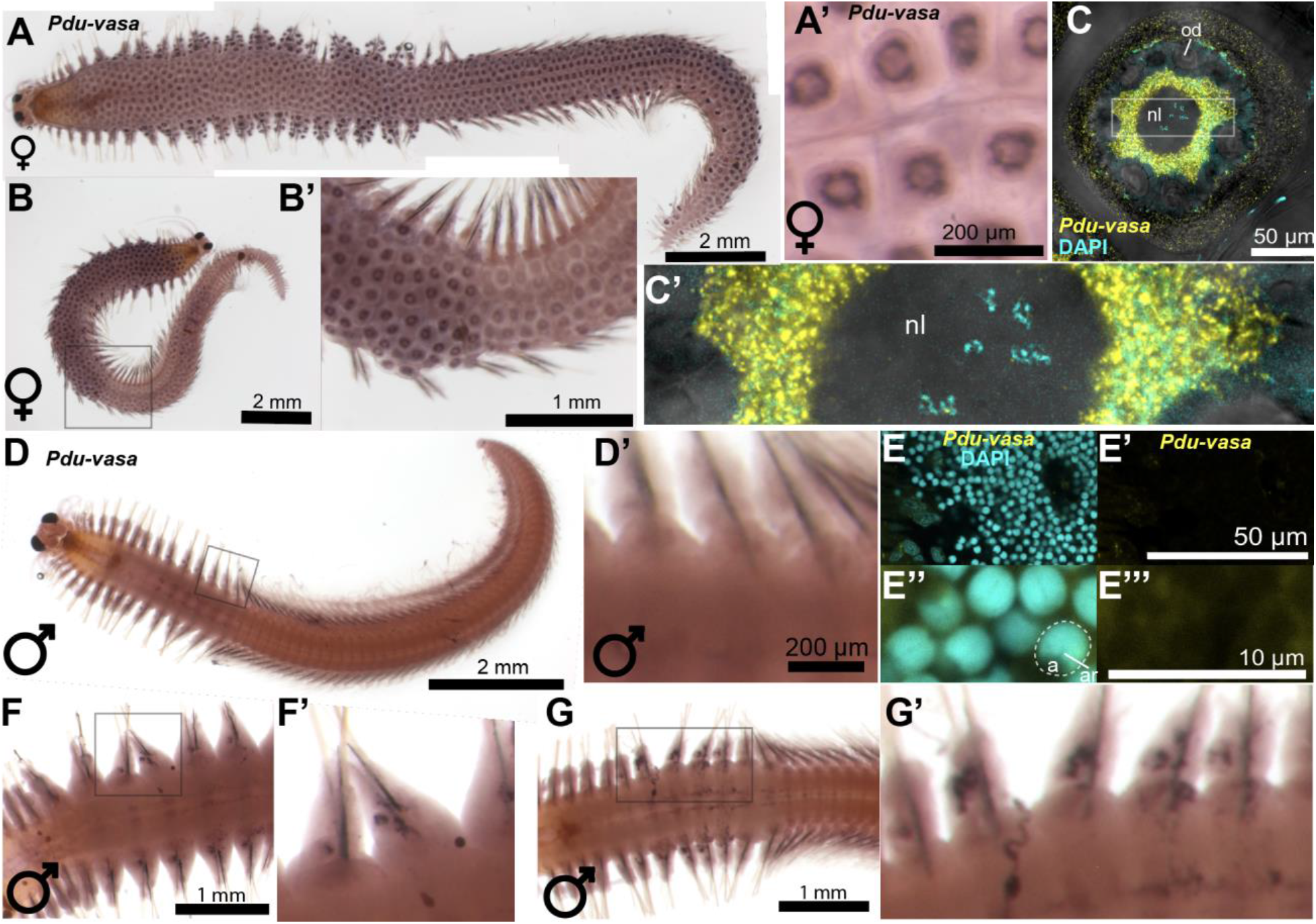
*vasa* expression in the mature female and males. **A-B’)** Sexually mature female vasa ISH shows vasa expression surrounding the nuclei of oocytes. **C-C’)** Fully mature oocyte processed for HCR showing *vasa* mRNA expression (yellow) and nuclear stain DAPI (cyan) in the perinuclear region. **D-D’)** Sexually mature male WISH shows no expression of *vasa*. However, depending on the level of maturation, some males still showed small patches of vasa positive regions **(F-G’). E-E’’’)** Spermatozoa in a fully mature male processed for HCR show no *vasa* signal. nl: nucleus, od: oil droplet, a: acrosome, ar: axial rod.

**Figure 4.**
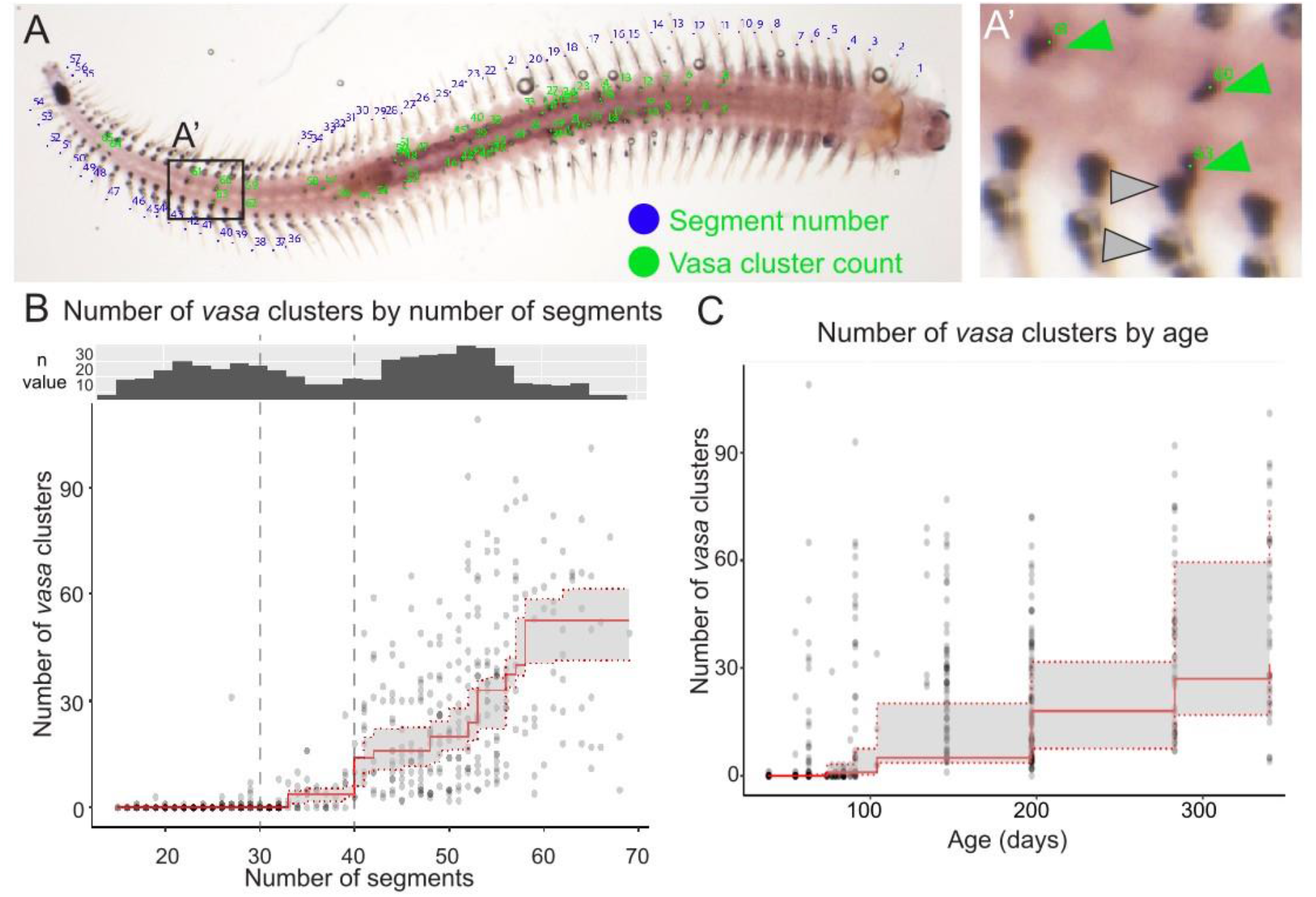
**A-A’)** Example scoring. A’) Gray arrowheads - background, not scored; Green arrowheads - *vasa*+ clusters, scored. **B)** Scatter plot showing the number of vasa clusters in worms by segment length at the time of fixation. Clusters typically begin forming after 30 segments and become numerous after approximately 40 segments. The estimated conditional median (red solid line) shows that this upward trend begins somewhere between 30-40 segments, closer to 35 segments. We also noted that there are no worms that are more than 40 segments in length that have zero vasa clusters in the trunk. The histogram above the scatter plot indicates the distribution of the number of segments in our dataset; notably, approximately half of the scored samples are below 40 segments and half are above. A 95% confidence interval for the conditional median is also indicated (gray shading between dashed red lines). **C)** The same data set and isotonic regression depicted in B but with median number of vasa clusters regressed on age. In contrast to segment number, age does not appear to be a strong determining factor for *vasa* cluster formation.

*P. dumerilii* has separate sexes. Towards completion of sexual maturation, females and males obtain morphological characteristics and secondary sexual structures that make it easy to differentiate them. However, this is not possible to do in the juveniles, which have very similar appearance in both sexes. We fixed female and male worms that started to mature sexually, carried out traditional colorimetric *in situ* hybridization or hybridization chain reaction (HCR, a fluorescent mRNA detection procedure) for *vasa*. In samples processed for colorimetric *in situ* hybridization, 22 out of 34 female samples exhibited strong *vasa* signal in oocytes with *vasa* signal localized perinuclear in the oocytes (Fig. 3A-C’). We confirmed this pattern of expression in the HCR-processed samples (Fig. 3C,C’). Among the colorimetric *in situ* hybridization samples, there were also some females without any signal (n=12/34). While this may be due to a technical factor such as probe penetration issues, these samples may also reflect a stage where detecting and/or visualizing vasa mRNA is harder because vasa mRNA may be too diffuse across the large oocyte volume. Indeed, we noticed gradations in the *vasa* signal that corresponded to how ready a female was to spawn based on external morphology and oocyte maturation stage. Females with morphology consistent to that of worms ready to spawn (n=22) (larger eggs, distinct atokous and epitokous parapodia) exhibited darker and perinuclear vasa signal compared to the females with less mature morphology (n=6) that lacked the distinct vasa signal (Suppl. Fig. 2). This is supported by HCR results where we observed the *vasa* signal in fully mature female oocytes of HCR-processed samples (n=2), but pre-mature females with oocytes still in maturation process (n=4) did not have the concentrated perinuclear vasa signal, and instead showed a much more diffuse signal across the oocyte (Suppl. Fig. 3). In most males, no signal for *vasa* expression was observed (Fig. 3D-G’), however, there were some males with spotty signal in their parapodia (n=13/54) (Fig. 3F-G’). We confirmed these results via HCR where no vasa signal was observed in the spermatozoa (Fig. 3E-E’’’). Sense controls showed no signal in oocytes or sperm (Suppl. Fig. 4B,C).

### *vasa*+ gonial cluster pattern is correlated with the total number of segments of individual worms

We next determined whether the chronological age (time since birth) or growth state (measured as total number of chaetigerous segments) of a worm had a greater effect on the progression of the pattern of *vasa*-expressing cells in the trunk. Specifically, we wanted to investigate when the switch from no vasa clusters to numerous vasa clusters happened. We hypothesized that if chronological age was the determining factor, worms would begin to produce *vasa*+ gonial clusters in the trunk at a set age regardless of size (in terms of segment number). If, however, chronological age was not critical, while reaching a particular growth state was the inducing factor for the vasa pattern, then we expected to see fast-growing worms develop vasa clusters more quickly relative to smaller worms of the same age.

To do this, we quantified the number of *vasa*+ gonial clusters in the trunk and the total number of segments for individual worms (n=494) (Fig. 4). Gonial cluster counts excluded the anterior cluster (see methods, Fig. 4A,A’). We also had chronological age information for all of these samples. Plotting *vasa*+ gonial clusters by number of segments showed clear correlation (Fig. 4B) while plotting *vasa*+ gonial clusters by age do not show a trend (Fig. 4C). Worms start forming *vasa*+ gonial clusters in the trunk when they reach 30-40 segments. However, during this growth stage, there are still numerous worms without any *vasa*+ gonial clusters in their trunk (n=41). In contrast, worms longer than 40 segments always have *vasa*+ gonial clusters in the trunk region, and the clusters become numerous after this segment length (Fig. 4B). Overall, these data suggest that worms show the *vasa*+ gonial cluster pattern based on the total segment number (growth) they have, and not based on their age (elapsed time since birth).

### Changing culture density slowed down growth and delayed *vasa*+ gonial cluster formation

We next tested whether culture density had any effect on the growth and gonial cluster formation by culturing worms in crowded or uncrowded conditions (100 worms in high density cultures and 20 worms in low density cultures). These cultures received the same amount of food per worm, were hosted in identical culture boxes, and overall kept under the same conditions except for the density of individuals in each box. We set the cultures up by splitting sibling worms into multiple high- or low-density cultures at 6 weeks post-fertilization (see Methods for details). Then, worms were cultured under these conditions for 2 weeks or 4 weeks, fixed and processed for whole mount *in situ* hybridization to investigate *vasa*+ gonial clusters (Fig. 5A). In addition, some worms from the same batch at the start of the experiment were fixed (t=0) (Fig. 5A, B).

**Figure 5.**
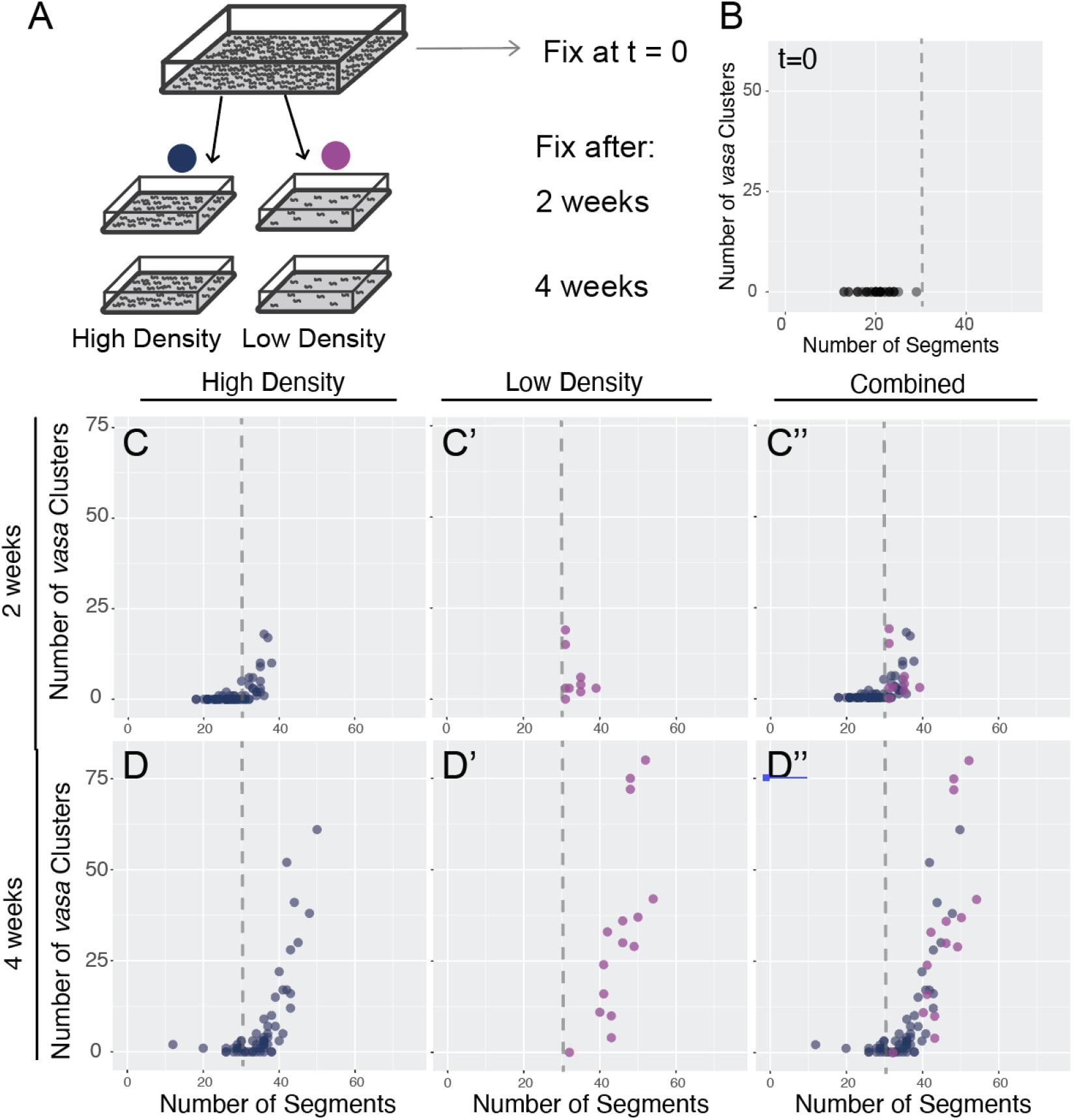
**A)** Experiment schematic showing the establishment of four new culture boxes from one large high density culture at six weeks post-fertilization. Two low density (n=20) and two high density cultures (n=100) were created. One of each culture type was fixed at two and four weeks post-establishment. B) Scatterplot showing number of vasa clusters and the number of segments per worm (n=39) taken at 6 weeks post-fertilization. C-C’’) Scatterplots showing the number of vasa clusters and the number of segments per worm two weeks after new cultures were established. D-D’’) Scatterplots showing the number of vasa clusters and the number of segments per worm four weeks after new cultures were established. Dashed line denotes 30 segment-length.

We found that in 2 weeks, all the worms in the low-density condition had developed several gonial clusters and were longer than 35 segments (Fig. 5C’) while only a fraction of the high-density culture samples had this phenotype (Fig. 5C). By 4 weeks under these conditions, most of the low-density samples had passed 40 segment length and developed numerous gonial clusters (Fig. 5D’) while the high density culture samples still had a larger fraction of worms below 40 segments (Fig. 5D). We repeated this experiment using ½ food per worm and found a similar result (Suppl. Fig. 5). Overall, these results suggest that despite having the same amount of food available, worms in high density conditions grow slower than the worms in low density conditions. However, irrespective of culture density conditions, the gonial cluster formation still follows a similar pattern: when worms reach 35-40 segments the number of gonial clusters increase, and almost all the worms longer than 40 segments have gonial clusters in their trunk segments. Therefore, while culture density affects the rate of growth, it does not affect the body size threshold effect on the progression of gametogenesis.

### Removing segments via amputation delays the *vasa*+ gonial cluster formation

Keeping worms in high density cultures slowed down growth, thus allowing us to examine the relationship of size to the production of the *vasa*+ gonial clusters. However, this experiment did not differentiate between whether worms were counting the total number of segments they generated since birth or whether reaching a particular growth state (i.e. minimum body size measured by number of segments) is enough for triggering the phenotype of having abundant *vasa*+ gonial clusters. To test this, we performed a series of tail amputation experiments to remove segments (i.e. to make shorter worms) and have worms generate new segments, while simultaneously keeping them below a certain segment number (Fig. 6A). These worms were the same age and were kept under the same culturing conditions but were reduced in size via amputation.

**Figure 6.**
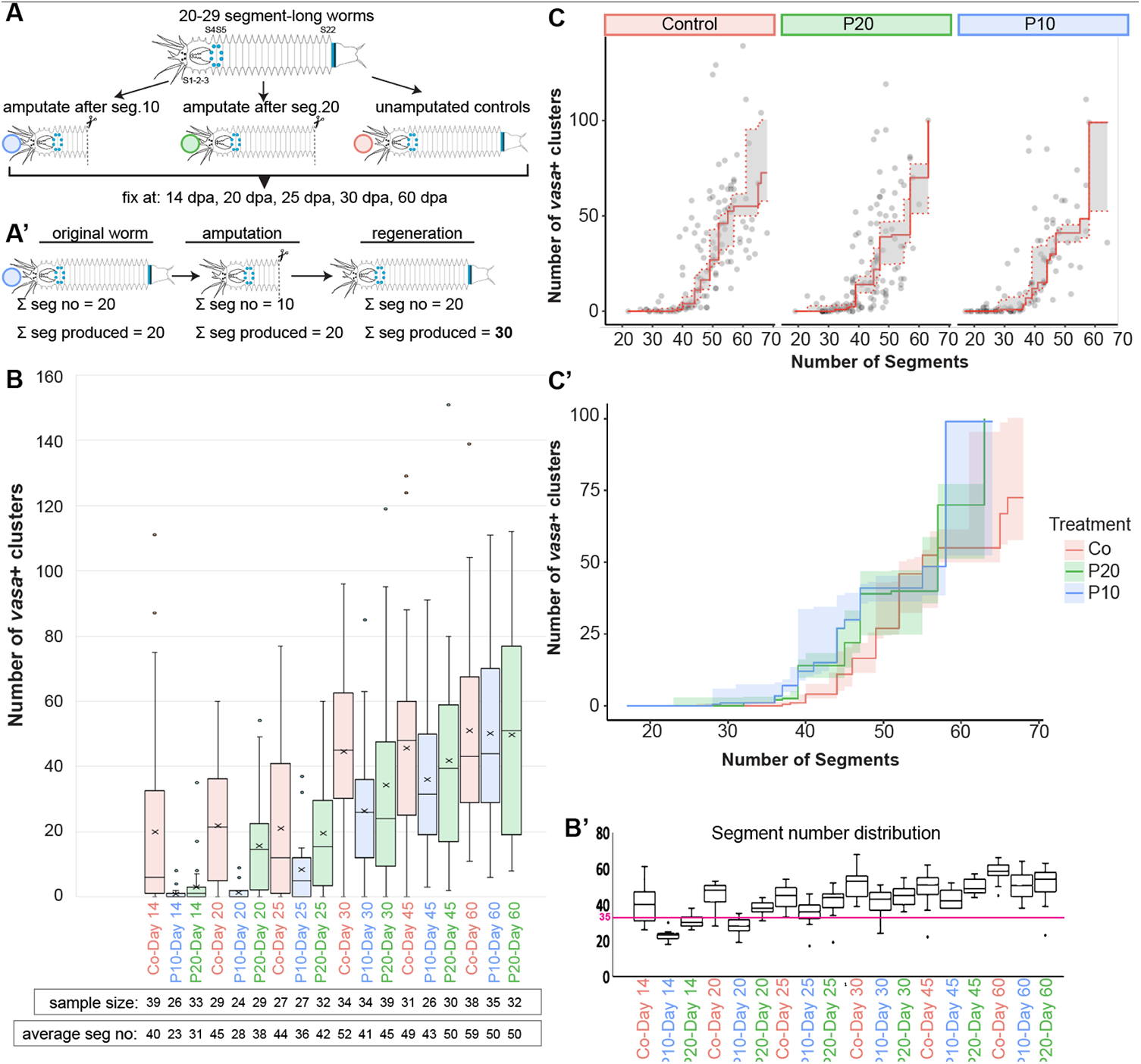
**A)** Amputation experiment design featuring controls, worms cut after segment 10, and worms cut after segment 20. All groups were fixed at the listed time points. A’) Examp Current length vs Total segment number generated during lifetime (including removed segments via amputation) **B)** Graph showing proportion of worms in each group that have 4 or more vasa positive clusters in their body cavity. **B’)** Segment number distribution for each group is shown. Magenta line denotes 35 segments, the size around which worms start showing an upward trend in forming vasa clusters. **C-C’)** Conditional median regression curves showing vasa cluster formation trends in each group, ignoring time, with 95% pointwise confidence intervals given by shaded regions (C’). All three lines show an upward trend around 35 segment length, which suggests the worms need to reach this absolute size threshold for forming vasa gonial clusters.

We began the experiment with worms that were 10-20 segments long, so that we could reasonably assume that they did not yet have any *vasa*+ gonial clusters in the trunk (Fig. 4B). We also fixed controls at the start (day 0) of the experiment and verified that this was true for this batch of worms (Dataset 03). Worms were separated into three groups: controls, worms cut after segment #10 (P10), and worms cut after segment #20 (P20). 30 worms in each group were fixed at different time points post-amputation (Fig. 6A), processed for whole mount ISH for *vasa* expression, and were scored for the number of *vasa*+ gonial clusters as described previously. The two amputated groups (P10 and P20) also served as controls for each other in case amputation alone caused an effect. Worms fully regenerate and produce approximately 10 new segments by 14 days post-amputation (Fig. 6B’).

If the worms are “counting” the total number of segments they have made and trigger *vasa*+ gonial cluster production accordingly, we would expect to see a shift in the appearance of gonial clusters to lower segment numbers. For example, if a worm of original length at 22 segments is amputated after segment 10, and regenerates 10 segments, it would have produced about 32 segments throughout its life (even though its current length would be 22 segments) (Fig. 6A’). Therefore, we would expect it to have a shift towards 20-30 segment length as the threshold for vasa cluster production, in contrast with 30-40 segment length in the control worms. However, if a minimum segment number at a given time (body size) is required for triggering the vasa cluster production, then we would expect to see a delay in amputated groups for producing vasa clusters, but the threshold would stay the same (30-40 segments).

We found that amputated worms in both groups P10 and P20 showed a delay in developing gonial clusters compared with controls (Fig. 6B). However, group P10 was more delayed than P20: for example, the majority of P20 samples fixed at day 20 of the experiment already started having gonial clusters while P10 samples at this time point were still mainly lacking vasa clusters (Fig. 6B). In all groups, as the worms get to the 30-40 segment threshold, they start forming gonial clusters (Fig. 6B,B’). To verify that the two experimental groups follow a similar trend compared with controls in terms for number of segments by the number of vasa clusters, we carried out isotonic median regression analyses (Fig. 6C, C’). We found that when we remove the time component, the vasa by segment curves are similar in all 3 groups: the trend curves show that once the experimental groups reach the similar segment numbers as controls, they have similar vasa cluster numbers. Therefore, the relationship between number of segments and (median) number of vasa clusters is not meaningfully different in each group. Overall, these results suggest that total segment number at a given time is the driving factor for the developmental stage transition, not the total number of segments a worm produced up to a certain time point.

### *vasa*+ gonial clusters are not required for regeneration

Some organisms such as hydra, planarians, or acoels have pluripotent stem cells scattered around the body and express germline/multipotency genes [33–35]. These cells are required for regeneration in these organisms. Recognizing that it is possible for a subset of the *vasa*+ cells in *Platynereis* could be similar to pluripotent stem cells in other highly regenerative organisms, we next asked if the clusters are required for regeneration in *Platynereis*. To address this, we analyzed the worms from the P10-P20 amputation experiment (Fig. 7). These worms are within the 10-20 segment length range and, as a result, do not have any *vasa*+ clusters in their trunk region, but only have the anterior cluster in the head region (around segments 4-5). Therefore, if the worms needed the trunk clusters to regenerate, they would fail regeneration. Alternatively, they would have cells from the anterior cluster respond to amputation and migrate towards the wound site. As expected, all the amputated worms regenerated, and we did not observe cells migrating towards the blastema from the anterior cluster region (Fig. 7B-D, I-K). In both groups, worms eventually formed *vasa*+ gonial clusters (Figs. 6 and 7F,G,M,N). Overall, these analyses show that neither the anterior *vasa*+ cells nor the trunk *vasa*+ clusters are required for regeneration in *Platynereis*.

**Figure 7.**
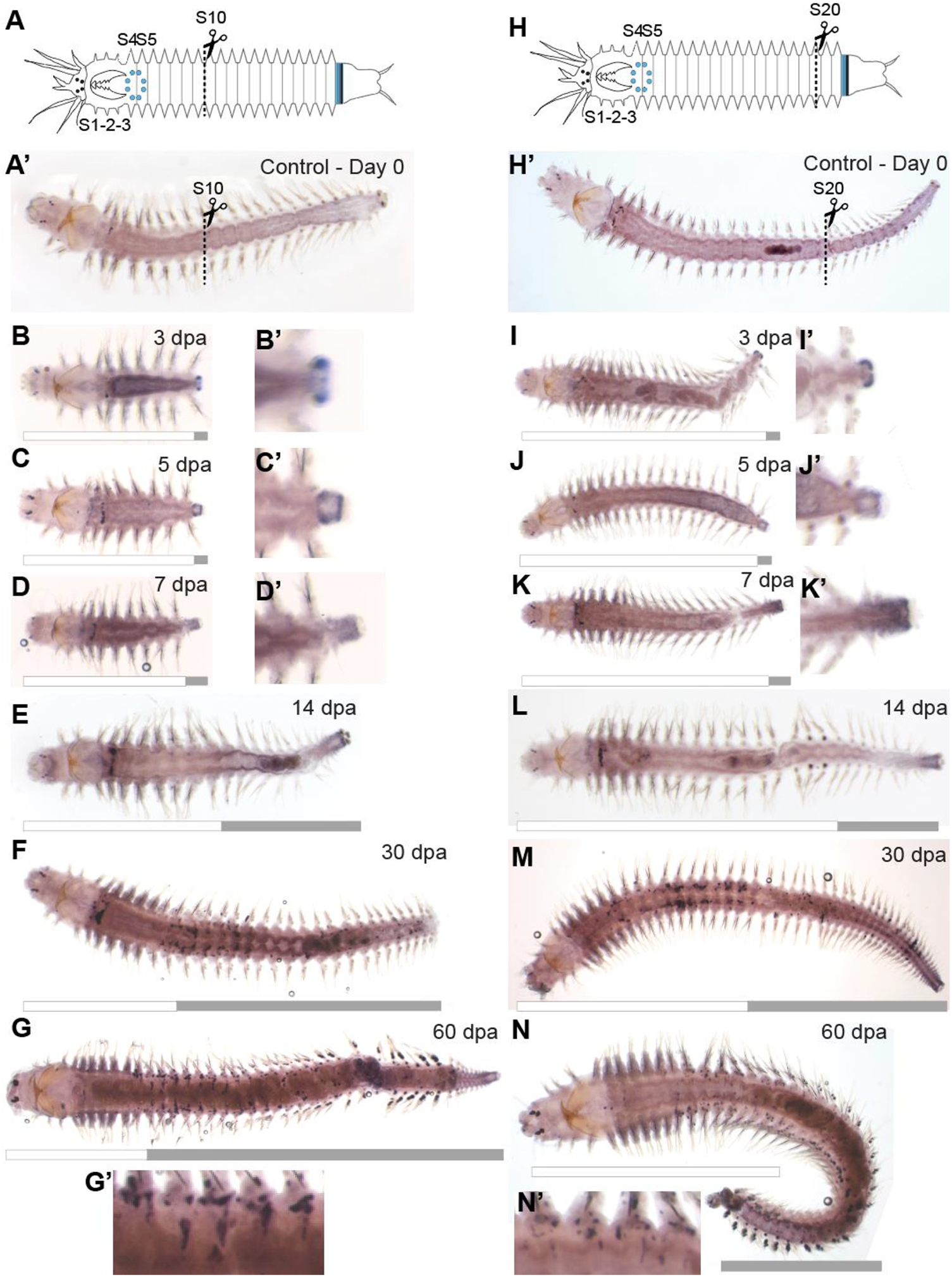
*vasa*+ cell clusters are not required for regeneration. A-A’ and H-H’ show amputation schematics and the control worms at the start of the experiment. B-G’) Worms amputated after segment 10, fixed at different time points and processed for WISH for *vasa*. I-N’) Worms amputated after segment 20, fixed at different time points and processed for WISH for *vasa*. Up until 30 dpa time point, neither group has vasa+ clusters in the trunk region other than the anterior-most cluster. However, all the samples regenerate successfully by forming a regeneration blastema which expresses vasa (B’-D’ and I’-K’). Gray bar denotes the regenerated region, while white bar denotes the original segments.

## DISCUSSION

### Gametogenesis in Platynereis dumerilii

Gametogenesis in Nereididae including *Platynereis dumerilii* has been extensively studied in juvenile worms that already have maturing oocytes that are 40 um or more in diameter or gonial clusters that already show signs of spermatogenesis [21,36–40]. These stages typically correspond to worms that are longer than 50 segments [41]. Stages of oocyte and sperm maturation in these larger worms have been studied using bright field and electron microscopy [15–17]. However, these studies predated the availability of molecular tools and therefore focused on stages of gametogenesis when it was more feasible to detect gonial clusters reliably (i.e. in worms longer than ~50 segments with maturing gametes). With the availability of *in situ* hybridization and immunohistochemistry, these works were followed by further analyses to investigate GMP marker expression in *P. dumerilii* across different stages of development including embryonic, larval, and juvenile worms [11–13]. However, a systematic analysis of formation of *vasa*+ gonial clusters in correlation with growth measured in segment number has not been carried out. Here, we addressed this knowledge gap and show that stages of gametogenesis in young juveniles are regulated by the growth state. Many previous studies in polychaetes used segment number as a metric, and this metric has been suggested as a staging system for juveniles in *P. dumerilii* previously as well [41]. Due to the wide variation in culture density and food availability across different studies, we suggest that chronological age is not a reliable method for staging juveniles. Our study reinforces that reporting the total segment number of juvenile *Platynereis* samples is more informative, if not necessary.

### Gametogenesis and growth

In many animals, the PGCs give rise to germline stem cells (GSCs), which in turn produce precursors that make the gametes (oocytes and sperm) [18,42]. These processes are often regulated by growth factors and hormones such as insulin. Nutritional status and insulin signaling affect GSC division rate in many organisms [4–6]. For example, *C. elegans* has germline specific insulin genes regulating germline proliferation [43]. Critical weight is required for metamorphosis in insects [7] and growth abnormalities delay developmental transitions in *Drosophila* [44]. Overall, growth, nutritional state, and sexual reproduction are intertwined [3,45,46].

The effects of hormones on growth, regeneration, and maturation have also been widely studied in *P. dumerilii* and other Nereididae [37,39,47–53]. However, these works either focus on larval stages, or the later stages of gametogenesis in premature worms that show signs of maturing gametes (see [47] for definitions of immature, premature, and mature). Our results suggest the presence of an earlier developmental stage transition (appearance of numerous gonial clusters) that is tightly coupled with growth. Whether there is hormonal regulation triggering *vasa*+ gonial cluster formation in these younger immature juveniles is still unknown. It will be interesting to investigate if estradiol, methylfarnesoate, and gonadotropin have a role during this developmental transition in the juveniles, as these two hormones have been studied at other developmental stages in *P. dumerilii* [48,49,51,52]. For example, homozygous crz1/gnrhl1 knockout animals show a delay in maturation. While this delay could be explained by reduced growth phenotype these mutants have, it would be curious to test if the mutants also have delayed formation of gonial clusters even when they reach the 35-40 segment size threshold. This could point out a decoupling of growth and gonial cluster formation. Similarly, methylfarnesoate, a member of the juvenile hormone family, has been identified as the brain hormone which has a role in regulating reproductive maturation [48]. Overall, expanding studies of gene and protein profiling [47], and testing the effects of different hormones including within the context of ecotoxicology at this juvenile developmental transition stage will reveal whether these hormones play a role in the developmental switch to start proliferating gonial clusters.

### Effect of culture density on growth and gonial cluster formation

Organisms in juvenile stages tend to grow in size as they age and progress towards sexual maturation. Therefore, in individually-reared organisms while age can be an acceptable predictor and descriptor of growth state, for organisms that are reared in batch cultures, the population density (or stocking density) can often have an effect on growth due to many factors such as accumulation of waste products, competition for food, and release of inhibiting substances by the organisms themselves [54–56]. Indeed, in our batch cultures, we often observe slower growth when worms are kept at high stocking density. We used these stocking density conditions as a way to slow down growth. We eliminated the age factor by using worms from the same clutch, and we reared them in different stocking density conditions to test for the effect of growth rate on the vasa+ gonial cluster numbers in individual worms. Worms kept in high stocking density grew slower (they had a slower rate of segment addition) than the worms in low stocking density. The reasons for the difference in growth rate between the high and low stocking density groups remain unclear. In all these groups, food was standardized so each worm theoretically had access to the same amount of calories. However, in low density cultures, there was more space per worm compared to in high density cultures. Having the additional space may help worms to grow more freely without risk of social conflict, and it may also help minimize the effects of waste products. Irrespective of the reasons for slowed growth, once the worms in high stocking density reached the threshold segment sizes (35-40 segments), they nevertheless formed *vasa*+ clusters similar to the low stocking density worms, as well as similar to the control worms in previous experiments. Therefore, while it took them longer to get to the threshold segment sizes, once they reached the threshold, they displayed similar patterns of gametogenesis. Overall, these results support the segment threshold idea and show that we did not elicit a substantial change in gonial cluster formation patterns in relation with growth state by changing the stocking density condition.

### Gametogenesis and “counting” segment number

Many annelids regulate and “count” segments they produce, although mechanisms for how they achieve this remain unclear. Polychaetes with indirect development typically have a species-specific number of segments produced in the larvae before hatching [57,58], and only after feeding and growth cues, they start producing new segments. In the clitellate annelid *Tubifex*, segment progenitor cells produced during embryogenesis have a birth rank order, and if these progenitors are transplanted in place of “earlier” ranks, they retain their birth rank, suggesting an internal “count” [59]. Therefore, in *P. dumerilii* we have addressed whether the worms were “counting” the number of segments they have produced, or whether they had to reach an absolute minimum size threshold. We assumed that if *P. dumerilii* juveniles were “counting” the total number of segments produced, with each new segment produced the worms could be growing their head region. This in turn could be triggering the transition into a developmental stage without needing to reach a minimum threshold body size. In other words, it was possible the brain underwent changes with each segment produced, even though worms were overall shorter after amputation, therefore the brain could exert an effect towards progressing gametogenesis irrespective of segment number. The effect of hormones produced by the brain on growth and reproduction has been well-studied [36,39,48,49,60]. However, our results support the minimum segment threshold hypothesis as the amputated worms still needed to reach 35-40 segment length for starting to form numerous gonial clusters. Of note, the amputated worms did not experience major differences in their rate of growth compared to controls, but there was a delay in germline development in amputated groups, the longest delays experienced by worms having undergone more severe amputations (P10). The increased lag in the P10 group, which had the more severe amputation, in comparison to the P20 group supports our claim from previous experiments that worms must reach a certain size before the gonial clusters can proliferate.

Unlike ecdysozoans, which have step function of growth, annelids have continuous asymptotic growth due to continuous segment addition [8]. Because the timing of sexual maturation can be predicted by reaching critical weight and molting, developmental transitions such as sexual maturation have been, arguably, easier to study in ecdysozoans. However, with organisms that show continuous growth such as annelids and planarians it is not as straightforward to predict the timing of a developmental transition based on body size/weight. While *P. dumerilii* is semelparous, and stops growing at maturation, during most of its lifetime, it shows continuous/indeterminate growth. Our work indicates an opportunity to address these questions using segment number as a reliable size/growth metric in *P. dumerilii*, which also has genetic and functional tools available as a research organism.

### Open questions about the gonial clusters and future directions

This study of systematic characterization of gonial cluster proliferation will help with developmental staging in juveniles for choosing the right individuals (based on segment number) to carry out experiments related to germline and gametogenesis. This will be specifically important for future studies of single cell transcriptomics analyses. Additionally, these results will help improve rearing conditions to maximize animal growth and maturation.

Many questions remain open, mainly on the sources of gonial clusters. Early embryonic origins of Primordial Germ Cells (PGCs) in *P. dumerilii* have been well investigated [11,14]. In the larvae, there is good evidence that the 4 PGCs migrate anteriorly behind the jaw region, and then proliferate in this location [11]. There is some evidence via DiI labeling and cell tracing that these cells from this anterior location (referred to as the anterior cluster in this manuscript) then migrate to more posterior trunk segments and form the gonial clusters in juvenile worms [12]. However, whether there are *vasa*+ gonial clusters originating from other sources such as locally in the trunk segments or from the the posterior growth zone, which is another region that expresses Germline/Multipotency (GMP) genes [24] such as *vasa*, remains to be addressed. In addition, there is good evidence that the *vasa*+ cell clusters are giving rise to gametes instead of being required for regeneration, but we cannot rule out the possibility that at least some of these cells may have different (non-germline) cell fates. Studies are underway using transgenesis to address these open questions in the future.

## ACKNOWLEDGEMENTS

Research reported in this publication was supported by the National Institute Of General Medical Sciences of the National Institutes of Health under Award Number R35GM138008 (to BDO) and R35GM133420 (to ADW) and Hibbitt Startup Funds (to BDO).

## COMPETING INTERESTS

Authors have no competing interests to declare.

## AUTHOR CONTRIBUTIONS

All authors have seen and approved the manuscript, and the manuscript has not been accepted or published elsewhere.

EK: experiments, experiment design, interpretation of results, data analysis, draft

BDO: experiment design, interpretation of results, data analysis, draft

DSC: data analysis and statistics

ADW: statistics

RWN: wrote the HCR oligo generator algorithm.

RWN and BM: carried out HCRs

## SUPPLEMENTARY MATERIAL

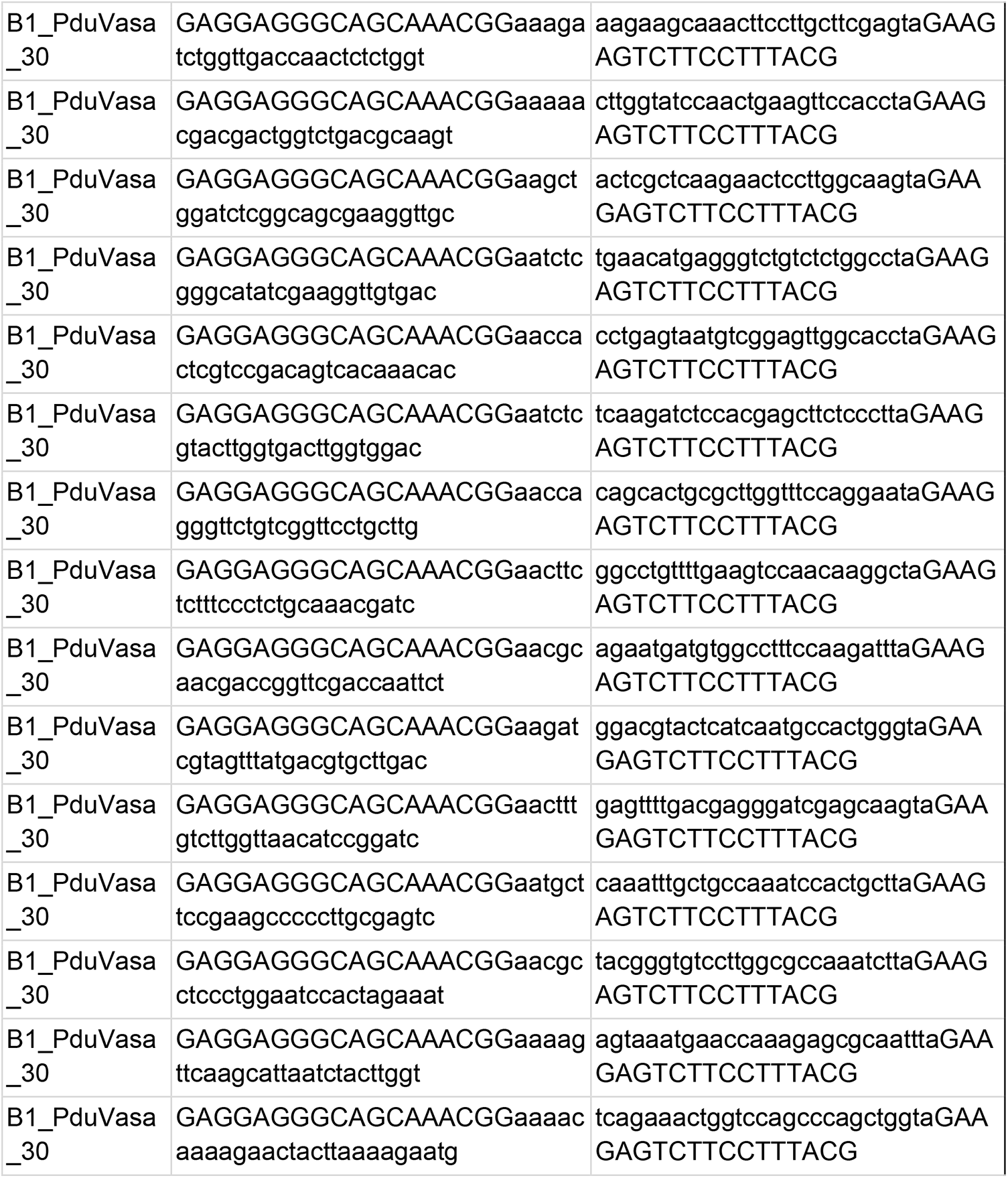
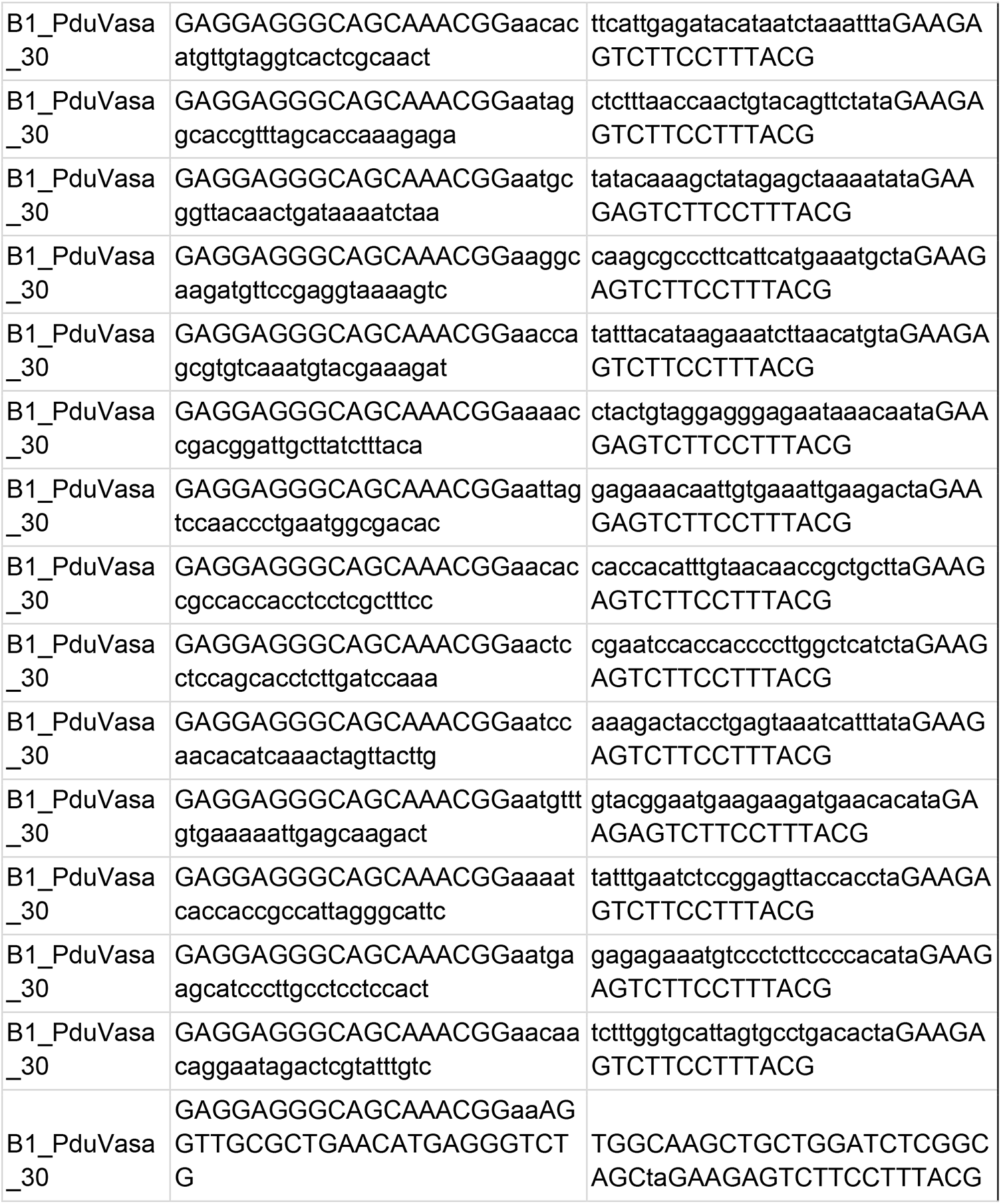
HCR Oligo pool for vasa mRNA detection. B1_PduVasa_30: Platynereis dumerilii vasa, with B1 amplifier, 30 pairs of oligos

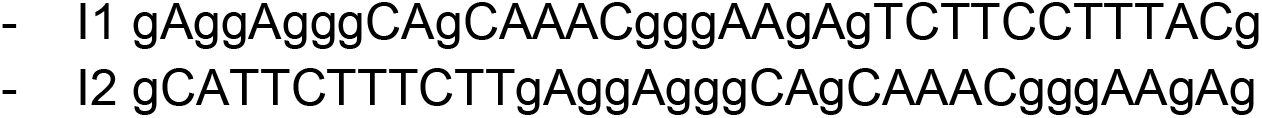
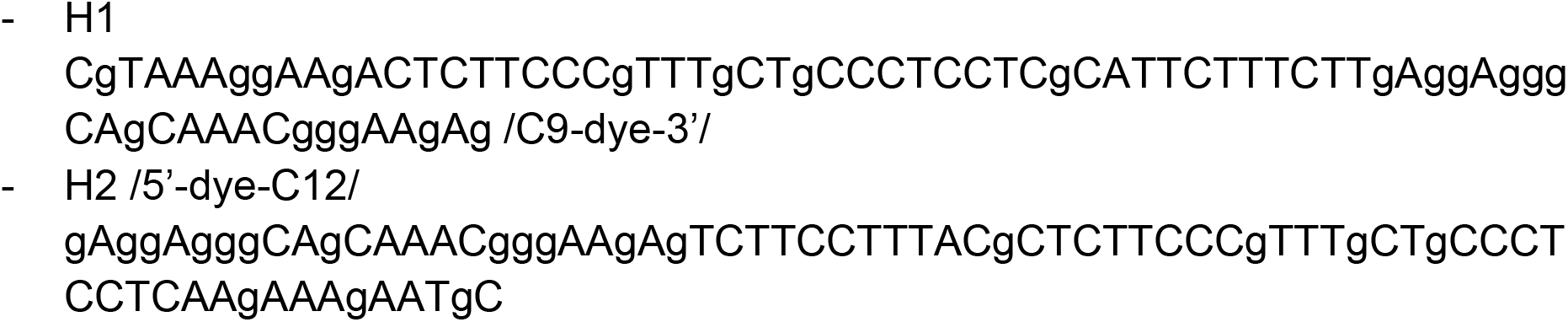
Hairpin Alexa-546 B1 from Molecular Instruments was used (Choi 2014)

## Food calculations for experiments

Normal feeding: 1.3 g food in 1 L FSW = 0.0013 g/mL

20 mL/box X 0.0013 g/mL - 0.026 g food/box

0.026 g / 30 worms = 0.00086 g food/worm

Fed twice/week: 0.00086 g x 2 = 0.00172 g/worm/week

Regular food (fed 3x per week): 0.00172 g / worm / week divided by 3 = 0.000573 g / worm = **0.57 mg / worm**

Half food (fed 3x per week): 0.000573 g / worm divided by 2 = 0.00028 g / worm = **0.28 mg / worm**

---

**Supplementary Figure 1.**
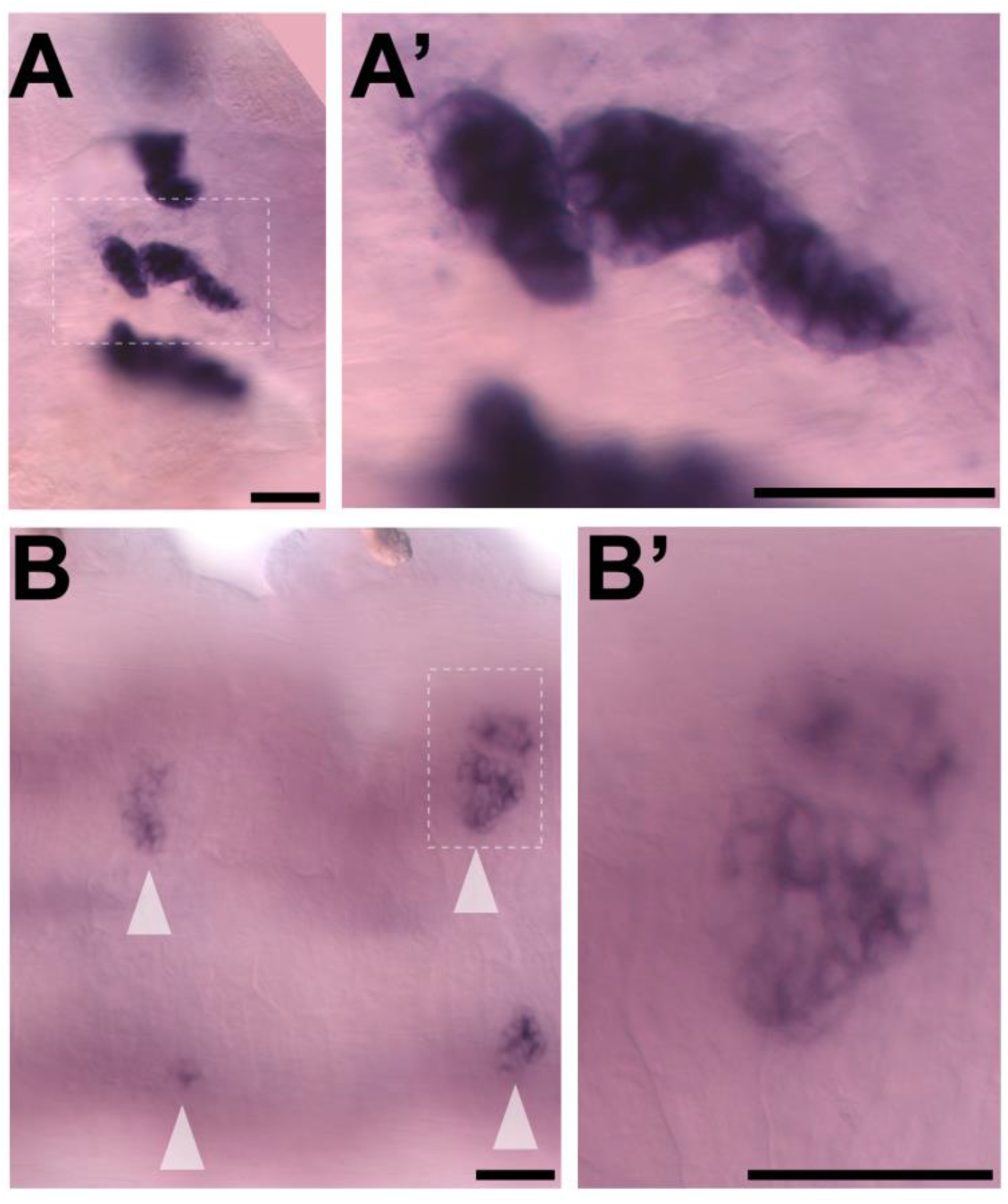
Close-up images of the anterior *vasa* cluster **(A-A’)** and trunk clusters **(B-B’)**. Samples processed for colorimetric *in situ* hybridization.

**Supplementary Figure 2.**
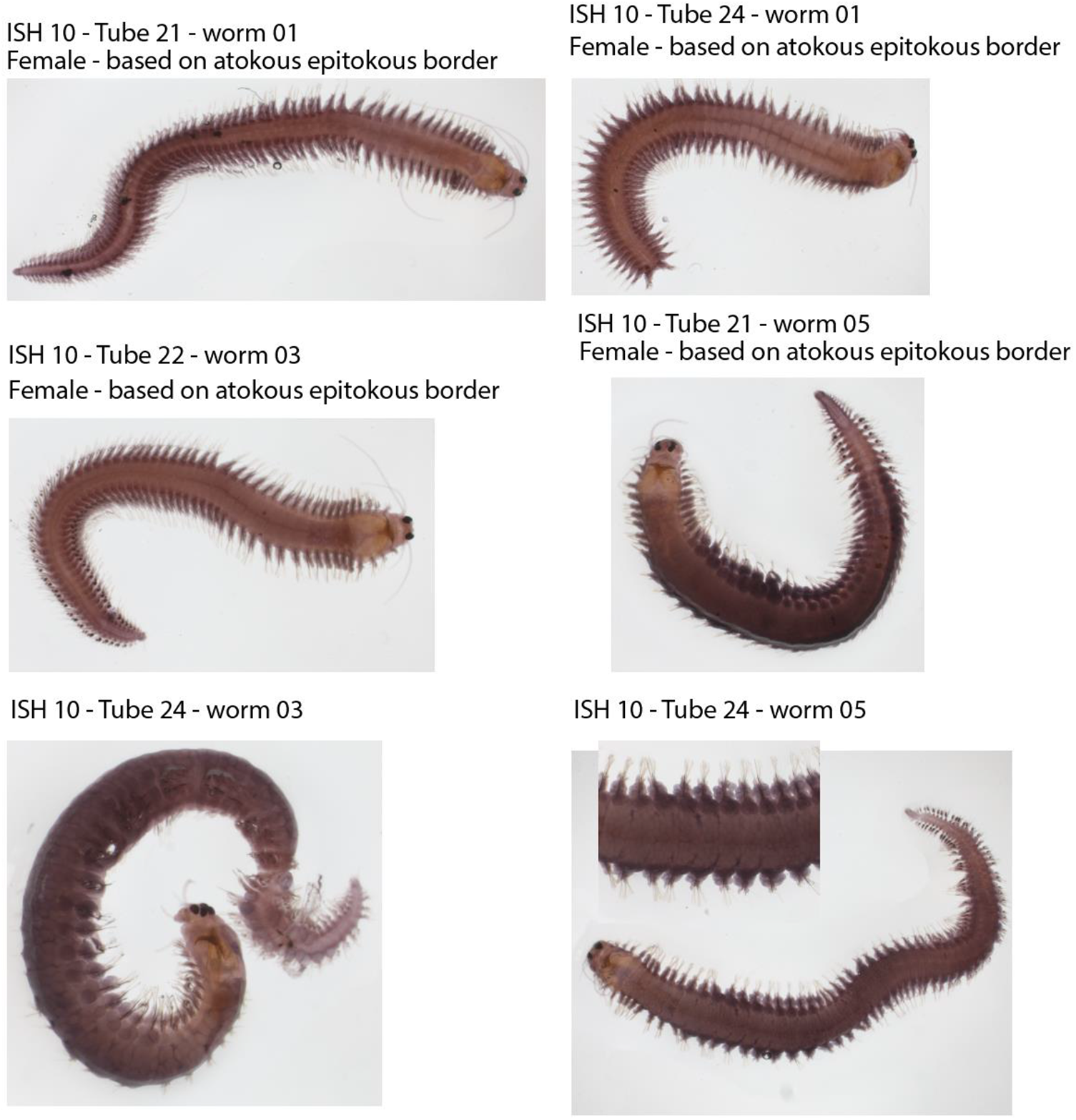
Examples of premature females, processed for WISH for *vasa* expression. The expression does not appear distinct and perinuclear as in fully mature females (Fig. 3).

**Supplementary Figure 3.**
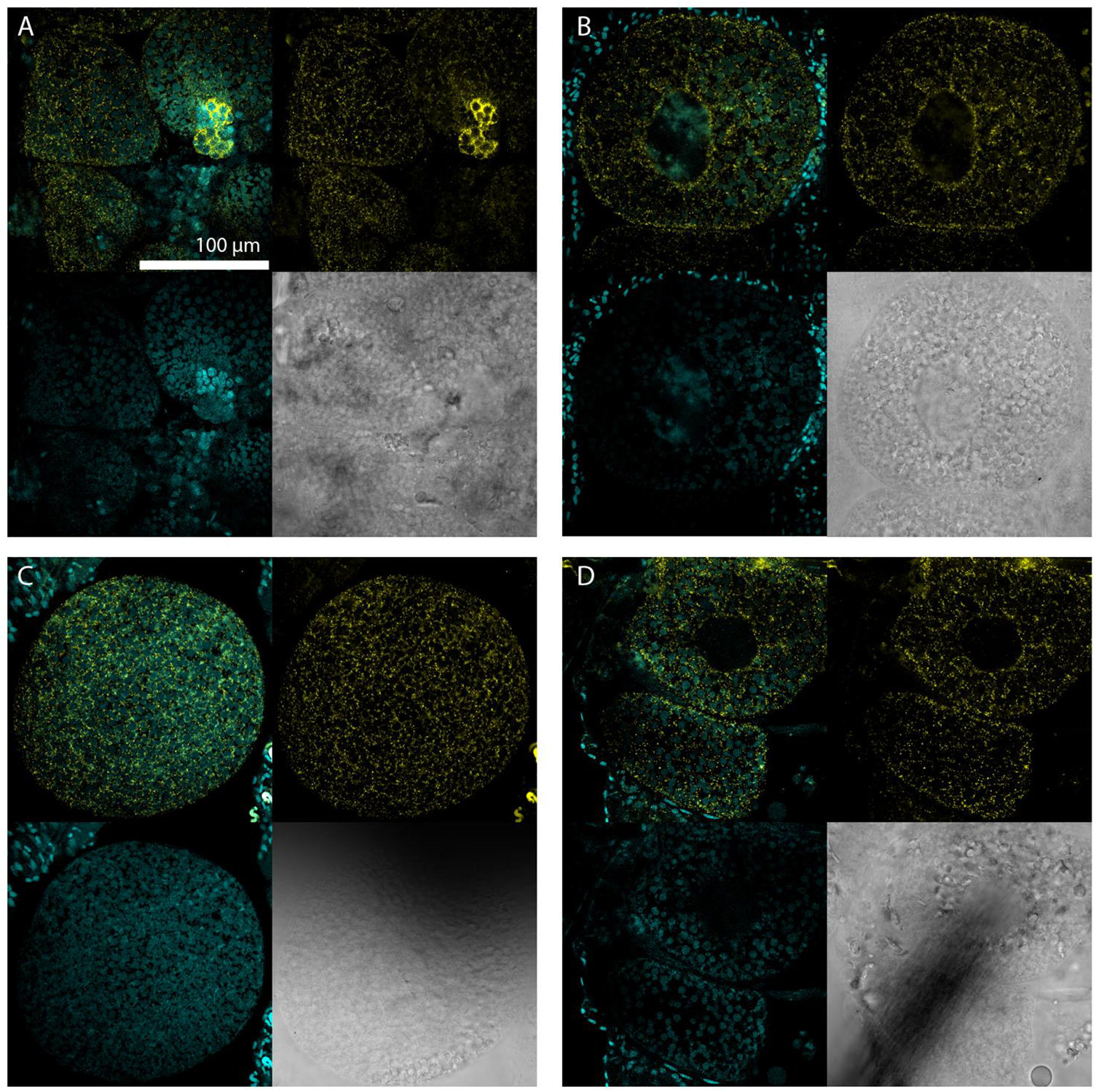
HCR for *vasa* mRNA in premature females (similar samples to Suppl. Fig. 2). Using this technique and confocal imaging, we were able to detect a diffuse signal and low level of *vasa* expression in the maturing oocytes. Note that there are smaller *vasa* clusters (in A) that are stained brightly, so penetration of probes and other reagents is not an issue. These oocytes appear to be still maturing because they lack the large and characteristic oil droplets, and distinct perinuclear region present in fully mature oocytes (Fig. 3C).

**Supplementary Figure 4.**
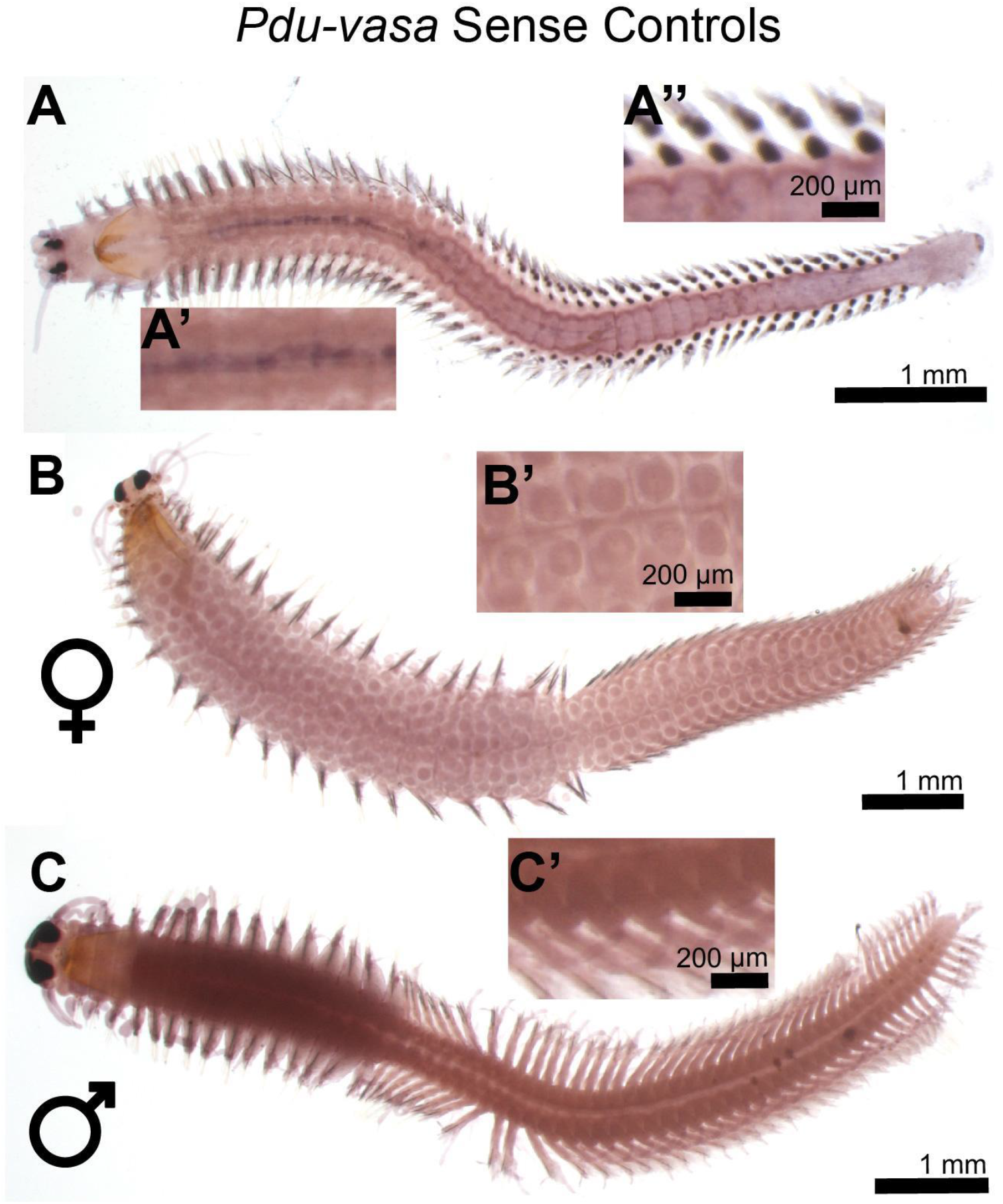
Sense control *in situ* hybridization for *vasa*. The sense probe was the same length as the antisense probe. **A)** Example of a juvenile worm with some background on the dorsal ectoderm **(A’)** and in chaetal sacs of parapodia **(A’’)**. **B)** Mature female. **C)** Mature male.

**Supplementary Figure 5.**
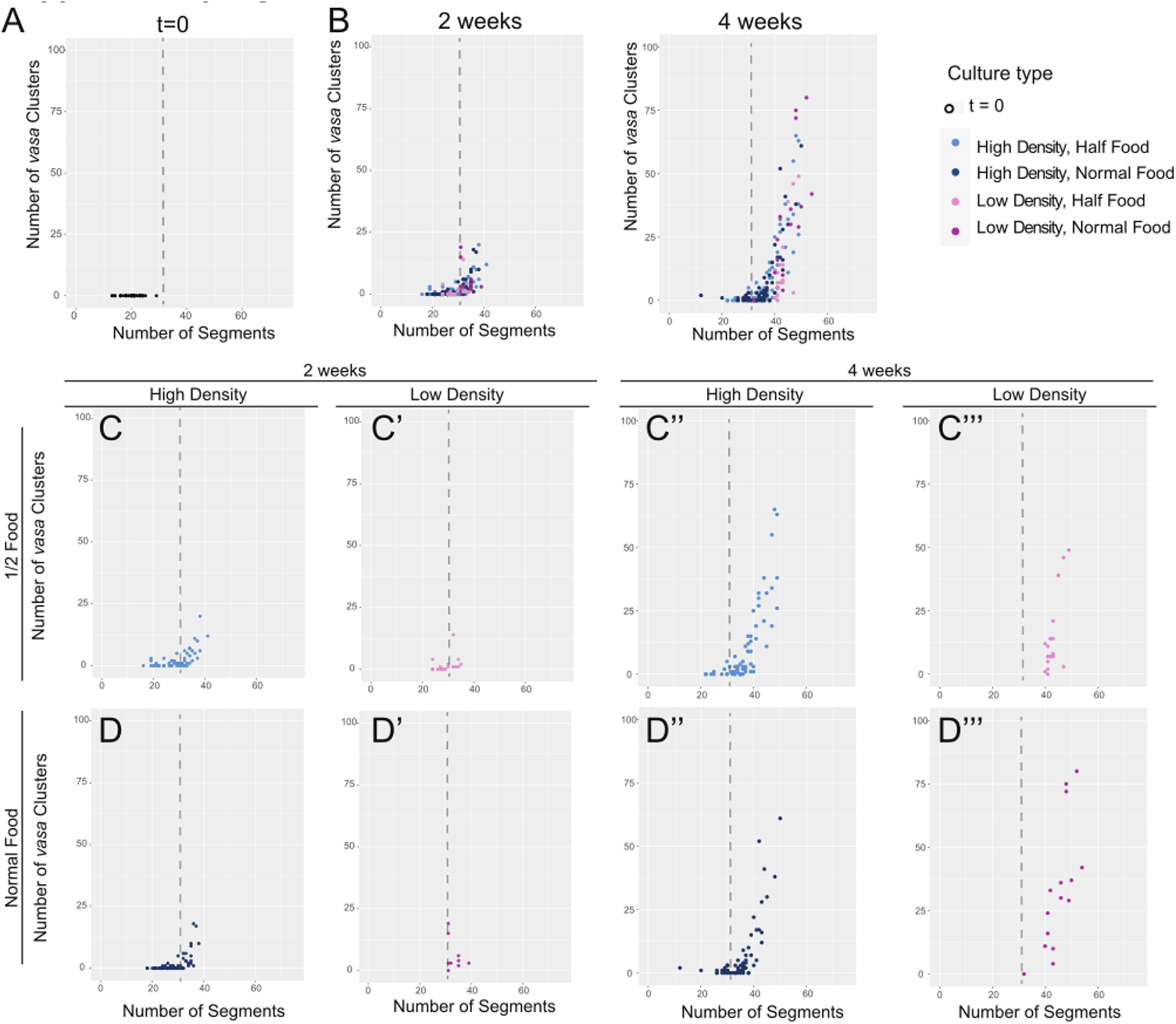
Scatter plots of half food vs normal food groups together. Same experiment as in Figure 5, and additional experiment with the same conditions except half the food was given to each box. A) Scatterplot showing number of vasa clusters and the number of segments per worm taken at 6 weeks post-fertilization at the start of the experiment. B) All data for experimental groups combined. C-C’’’) Individual scatterplots showing the number of vasa clusters and the number of segments per worm in the "half food" group per time point. D-D’’) Scatterplots showing the number of vasa clusters and the number of segments per worm in the "normal food" group (same data as in Fig. 5). Dashed line denotes 30 segment-length.

**Supplementary Figure 6.**
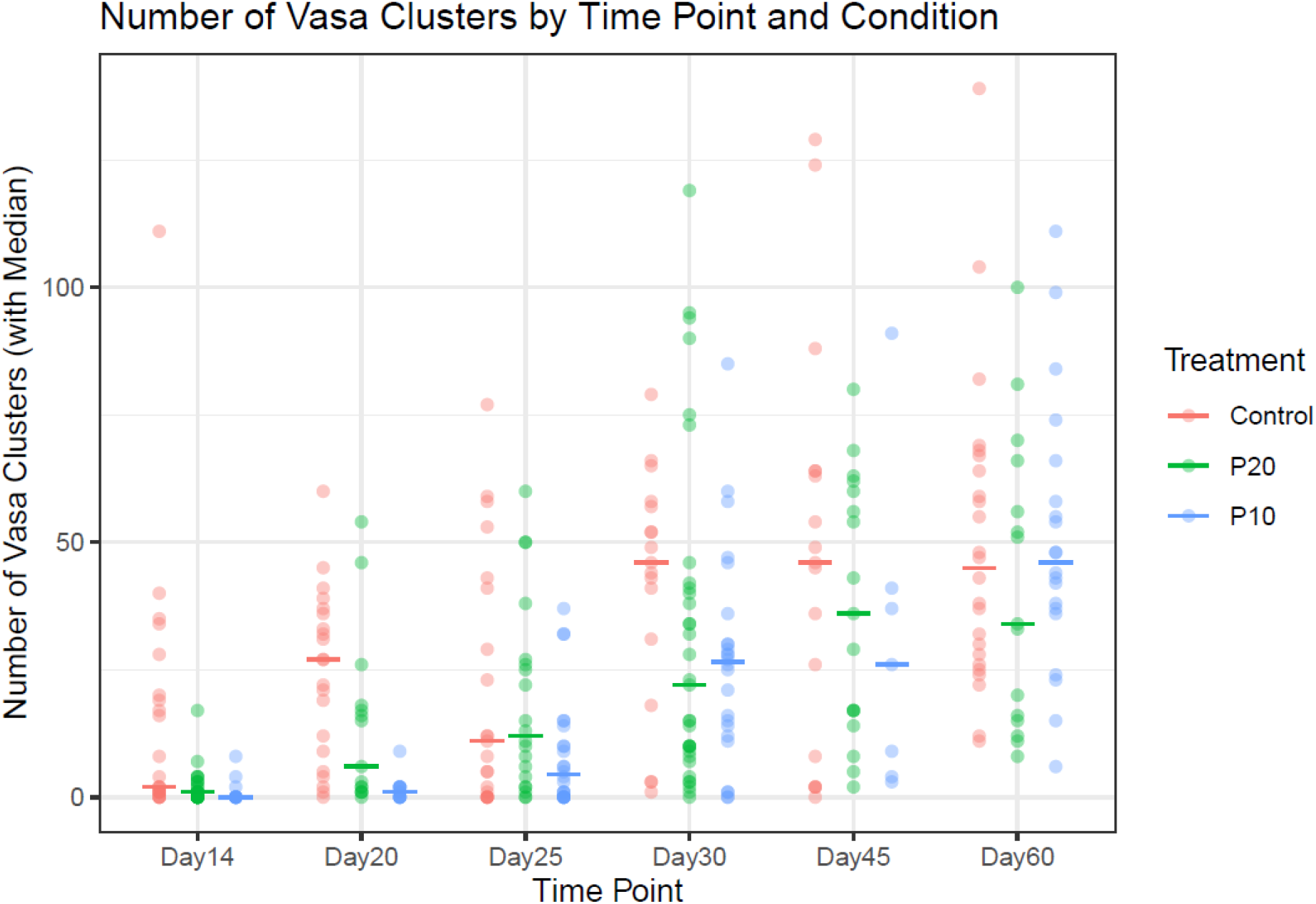
Another representation of the box plot in Fig. 6B. This plot shows 95% bootstrap confidence intervals (horizontal bars) for the median in each group. We note that a cluster bootstrap is not feasible here (each treatment-day combination consists of a single culture box), so these confidence intervals do not account for possible box effects.

## Notes

https://github.com/BDuyguOzpolat/Platynereis-gametogenesis-Kuehn-et-al-2021/blob/main/README.md

### Competing Interest Statement

The authors have declared no competing interest.

## REFERENCES

1. Lord JP, Shanks AL. 2012 Continuous growth facilitates feeding and reproduction: Impact of size on energy allocation patterns for organisms with indeterminate growth. 159, 1417–1428.

2. Lui JC, Baron J. 2011 Mechanisms limiting body growth in mammals. Endocr. Rev. 32, 422–440.

3. Hyun S. 2018 Body size regulation by maturation steroid hormones: a Drosophila perspective. Front. Zool. 15, 44.

4. Shim J, Gururaja-Rao S, Banerjee U. 2013 Nutritional regulation of stem and progenitor cells in Drosophila. Development 140, 4647–4656.

5. Narbonne P, Roy R. 2006 Regulation of germline stem cell proliferation downstream of nutrient sensing. Cell Div. 1, 29.

6. Hubbard EJA, Korta DZ, Dalfó D. 2013 Physiological control of germline development. Adv. Exp. Med. Biol. 757, 101–131.

7. Tennessen JM, Thummel CS. 2011 Coordinating growth and maturation - insights from Drosophila. Curr. Biol. 21, R750–7.

8. Sebens KP. 1987 The ecology of indeterminate growth in animals. Annu. Rev. Ecol. Syst. 18, 371–407.

9. Hutchinson JMC, McNamara JM, Houston AI, Vollrath F. 1997 Dyar’s Rule and the Investment Principle: optimal moulting strategies if feeding rate is size–dependent and growth is discontinuous. Philos. Trans. R. Soc. Lond. B Biol. Sci. 352, 113–138.

10. Rebscher N. 2014 Establishing the germline in spiralian embyos. Int. J. Dev. Biol. 58, 403–411.

11. Rebscher N, Lidke AK, Ackermann CF. 2012 Hidden in the crowd: primordial germ cells and somatic stem cells in the mesodermal posterior growth zone of the polychaete Platynereis dumerillii are two distinct cell populations. Evodevo 3, 9.

12. Rebscher N, Zelada-González F, Banisch TU, Raible F, Arendt D. 2007 Vasa unveils a common origin of germ cells and of somatic stem cells from the posterior growth zone in the polychaete Platynereis dumerilii. Dev. Biol. 306, 599–611.

13. Zelada González YF. 2005 Germline development in Platynereis dumerilii and its connection to embryonic patterning. (doi:10.11588/heidok.00005432)

14. Özpolat BD, Handberg-Thorsager M, Vervoort M, Balavoine G. 2017 Cell lineage and cell cycling analyses of the 4d micromere using live imaging in the marine annelid platynereis dumerilii. Elife 6. (doi:10.7554/eLife.30463)

15. Fischer A. 1975 The structure of symplasmic early oocytes and their enveloping sheath cells in the polychaete, Platynereis dumerilii. Cell Tissue Res. 160, 327–343.

16. Fischer A. 1974 Stages and stage distribution in early oogenesis in the Annelid, Platynereis dumerilii. Cell Tissue Res. 156, 35–45.

17. Meisel J. 1990 Zur Hormonabhängigkeit der Spermatogenese bei Platynereis dumerilii: licht-und elektronenmikroskopische Befunde sowie experimentelle Untersuchungen in vivo und in vitro. na.

18. Nieuwkoop PD, Sutasurya LA. 1981 Primordial Germ Cells in the Invertebrates: From Epigenesis to Preformation. Cambridge: Cambridge University Press.

19. Giese AC, Pearse J, editors. 1974 Reproduction of Marine Invertebrates - Acoelomate and Pseudocoelomate Metazoans. Academic Press.

20. Hempelmann F. 1911 Zur Naturgeschichte von Nereis dumerilii Aud. et Edw.

21. Hauenschild C, Fischer A. 1969 Platynereis dumerilii: mikroskopische Anatomie, Fortpflanzung, Entwicklung. Stuttgart: G. Fischer.

22. ÖzpolatLab-GitHub-Kuehn2021. In press. R codes, protocols, additional files on Github for Kuehn et al 2021. Github. See https://github.com/BDuyguOzpolat/Platynereis-gametogenesis-Kuehn-et-al-2021.

23. Kuehn E, Stockinger AW, Girard J, Raible F, Özpolat BD. 2019 A scalable culturing system for the marine annelid Platynereis dumerilii. PLoS One 14, e0226156.

24. Gazave E, Béhague J, Laplane L, Guillou A, Préau L, Demilly A, Balavoine G, Vervoort M. 2013 Posterior elongation in the annelid Platynereis dumerilii involves stem cells molecularly related to primordial germ cells. Dev. Biol. 382, 246–267.

25. Choi HMT, Beck VA, Pierce NA. 2014 Next-generation in situ hybridization chain reaction: higher gain, lower cost, greater durability. ACS Nano 8, 4284–4294.

26. Choi HMT, Schwarzkopf M, Fornace ME, Acharya A, Artavanis G, Stegmaier J, Cunha A, Pierce NA. 2018 Third-generation in situ hybridization chain reaction: multiplexed, quantitative, sensitive, versatile, robust. Development 145. (doi:10.1242/dev.165753)

27. ÖzpolatLab-HCR. In press. Özpolat Lab HCR probe generator. Github. See https://github.com/rwnull/insitu_probe_generator.

28. Schindelin J et al. 2012 Fiji: an open-source platform for biological-image analysis. Nat. Methods 9, 676–682.

29. RStudio Team. 2020 RStudio: Integrated Development Environment for R.

30. R Core Team. 2020 R: A Language and Environment for Statistical Computing.

31. de Leeuw J, Hornik K, Mair P. 2009 Isotone Optimization in R: Pool-Adjacent-Violators Algorithm (PAVA) and Active Set Methods. Journal of Statistical Software, Articles 32, 1–24.

32. Abrevaya J. 2005 Isotonic Quantile Regression: Asymptotics and Bootstrap. J. Indian Soc. Agric. Stat. 67, 187–199.

33. Gehrke AR, Srivastava M. 2016 Neoblasts and the evolution of whole-body regeneration. Curr. Opin. Genet. Dev. 40, 131–137.

34. David CN. 2012 Interstitial stem cells in Hydra: multipotency and decision-making. Int. J. Dev. Biol. 56, 489–497.

35. Shibata N, Umesono Y, Orii H, Sakurai T, Watanabe K, Agata K. 1999 Expression of vasa(vas)-related genes in germline cells and totipotent somatic stem cells of planarians. Dev. Biol. 206, 73–87.

36. Hauenschild C. 1974 Normalisierung der geschlechtlichen Entwicklung kopfloser Fragmente junger ♀♀ vonPlatynereis dumerilii (Polychaeta) durch Behandlung mit konservierten Prostomien juveniler Individuen. Helgoländer wissenschaftliche Meeresuntersuchungen 26, 63–81.

37. Lawrence AJ, Soame JM. 2009 The endocrine control of reproduction in Nereidae: a new multi-hormonal model with implications for their functional role in a changing environment. Philos. Trans. R. Soc. Lond. B Biol. Sci. 364, 3363–3376.

38. Brafield AE, Chapman G. 1967 Gametogenesis and Breeding in a Natural Population of Nereis Virens. J. Mar. Biol. Assoc. U. K. 47, 619–627.

39. Schroeder PC, Hermans CO. 1975 Annelida: Polychaeta. In Reproduction of Marine Invertebrates. Vol. 3: Annelids and Echiurans (eds AC Giese, JS Pearse), New York: Academic Press.

40. Fallon JF, Austin CR. 1967 Fine structure of gametes of Nereis limbata (Annelida) before and after interaction. J. Exp. Zool. 166, 225–241.

41. Fischer AH, Henrich T, Arendt D. 2010 The normal development of Platynereis dumerilii (Nereididae, Annelida). Front. Zool. 7, 31.

42. Nieuwkoop PD, Sutasurya LA. 1979 Primordial Germ Cells in the Chordates: Embryogenesis and Phylogenesis. Cambridge: Cambridge University Press.

43. Michaelson D, Korta DZ, Capua Y, Hubbard EJA. 2010 Insulin signaling promotes germline proliferation in C. elegans. Development 137, 671–680.

44. Hariharan IK. 2012 How growth abnormalities delay ‘puberty’ in Drosophila. Sci. Signal. 5, e27.

45. Tharp ME, Collins JJ 3rd, Newmark PA. 2014 A lophotrochozoan-specific nuclear hormone receptor is required for reproductive system development in the planarian. Dev. Biol. 396, 150–157.

46. Plaistow SJ, Lapsley CT, Beckerman AP, Benton TG. 2004 Age and size at maturity: sex, environmental variability and developmental thresholds. Proc. Biol. Sci. 271, 919–924.

47. Schenk S et al. 2019 Combined transcriptome and proteome profiling reveals specific molecular brain signatures for sex, maturation and circalunar clock phase. Elife 8. (doi:10.7554/eLife.41556)

48. Schenk S, Krauditsch C, Frühauf P, Gerner C, Raible F. 2016 Discovery of methylfarnesoate as the annelid brain hormone reveals an ancient role of sesquiterpenoids in reproduction. Elife 5. (doi:10.7554/eLife.17126)

49. Andreatta G et al. 2019 Corazonin signaling integrates energy homeostasis and lunar phase to regulate aspects of growth and sexual maturation in Platynereis. Proc. Natl. Acad. Sci. U. S. A. (doi:10.1073/pnas.1910262116)

50. Hofmann DK. 1976 Regeneration and endocrinology in the polychaetePlatynereis dumerilii. Wilhelm Roux’s Archives of Developmental Biology 180, 47–71.

51. García-Alonso J, Ayoola JAO, Crompton J, Rebscher N, Hardege JD. 2011 Development and maturation in the nereidid polychaetes Platynereis dumerilii and Nereis succinea exposed to xenoestrogens. Comp. Biochem. Physiol. C. Toxicol. Pharmacol. 154, 196–203.

52. Lidke AK et al. 2014 17β-Estradiol induces supernumerary primordial germ cells in embryos of the polychaete Platynereis dumerilii. Gen. Comp. Endocrinol. 196, 52–61.

53. Golding DW. 1967 ENDOCRINOLOGY, REGENERATION AND MATURATION IN NEREIS. Biol. Bull. 133, 567–577.

54. Tettelbach ST et al. 2011 Utility of high-density plantings in bay scallop, Argopecten irradians irradians, restoration. Aquac. Int. 19, 715–739.

55. Biswas JK, Sarkar D, Chakraborty P, Bhakta JN, Jana BB. 2006 Density dependent ambient ammonium as the key factor for optimization of stocking density of common carp in small holding tanks. Aquaculture 261, 952–959.

56. Dong J, Zhao Y-Y, Yu Y-H, Sun N, Li Y-D, Wei H, Yang Z-Q, Li X-D, Li L. 2018 Effect of stocking density on growth performance, digestive enzyme activities, and nonspecific immune parameters of Palaemonetes sinensis. Fish Shellfish Immunol. 73, 37–41.

57. Blake JA. 2017 Larval development of Polychaeta from the northern California coast. Fourteen additional species together with seasonality of planktic larvae over a 5-year period. J. Mar. Biol. Assoc. U. K. 97, 1081–1133.

58. Balavoine G. 2014 Segment formation in Annelids: patterns, processes and evolution. Int. J. Dev. Biol. 58, 469–483.

59. Kato Y, Nakamoto A, Shiomi I, Nakao H, Shimizu T. 2013 Primordial germ cells in an oligochaete annelid are specified according to the birth rank order in the mesodermal teloblast lineage. Dev. Biol. 379, 246–257.

60. Hauenschild C. 1966 Der hormonale einfluss des Gehirns auf die sexuelle Entwicklung bei dem polychaeten Platynereis dumerilii. Gen. Comp. Endocrinol. 6, 26–73.

